# The first OpenBind release: An open experimental structure–affinity dataset and benchmark for structure-based AI

**DOI:** 10.64898/2026.08.27.747600

**Authors:** Jochem Nelen, Omeir Khan, Etowah Adams, Jasmin C. Aschenbrenner, Warren Thompson, Ali Ebrahim, Eda Çapkin, Cédric Vallée, OpenBind, Elizabeth J. Shotton, Ed J. Griffen, John D. Chodera, Charlotte M. Deane, Frank von Delft, Mohammed AlQuraishi, Fergus Imrie

## Abstract

High-quality experimental datasets that link protein–ligand structures with binding affinity data are essential for developing and evaluating structure-based machine learning methods. To help address this need, we established OpenBind as an open-science initiative to generate large-scale experimental datasets for structure-based AI and molecular discovery. Here, we describe the first public OpenBind release, which, to the best of our knowledge, is the largest public single-target experimental structure–affinity dataset. The dataset focuses on enteroviral 2A protease, comprising 925 crystallographic binding events from 699 compounds and associated affinity measurements for 601 compounds. It combines structures from an initial fragment screen and follow-on molecules, together with affinity data, linking experimentally determined protein–ligand binding modes to biophysical measurements within a coherent antiviral discovery campaign. We used this dataset to evaluate protein–ligand structure prediction, binding-affinity prediction, and virtual screening using representative structure-based methods, including docking and cofolding. This exposed several challenges that are central to practical structure-based modelling: docking performance depends strongly on binding-pocket conformation, poses are difficult to rank, and structure-based affinity prediction remains challenging. Fine-tuning OpenFold3-p2 on the fragment-screen structures substantially improved pose prediction and virtual screening for related follow-on compounds, demonstrating how early-stage experimental structures can support target-specific model adaptation.

## 1 Introduction

Recent advances in deep learning have substantially expanded the scope of computational structural biology and structure-based drug discovery [1]. AlphaFold2 [2] demonstrated that machine learning (ML)-based approaches can achieve highly accurate protein structure prediction at the proteome scale, while subsequent open-source implementations such as OpenFold [3] also provided training code and data [4], enabling broader model development and adoption. The next generation of models, most notably AlphaFold3 [5], extended these ideas to biomolecular complex prediction and protein–ligand modelling, marking another advance beyond protein structure prediction alone.

However, accurate protein–ligand complex prediction remains challenging. Although recent cofolding methods have achieved relatively high accuracy for complexes resembling structures available during model development, their performance drops substantially for more novel protein–ligand systems [6]. Further, structure prediction is only part of the solution as structure-based drug discovery also requires reliable binding affinity estimates for effective compound prioritisation. This remains a difficult and largely unresolved problem, since binding affinity depends on a complex combination of factors that are challenging to model accurately, including subtle intermolecular interactions, protein and ligand flexibility, solvation effects, and entropic contributions [7]. Recent methods such as Boltz-2 [8], which jointly predicts protein–ligand structures and binding affinity, and AQAffinity [9], an affinity-prediction head built on OpenFold3, represent important attempts to address this gap. Nevertheless, recent evaluations suggest current cofolding-based affinity predictions are not yet sufficiently robust for reliable compound ranking [10], despite encouraging results in some affinity-ranking and hit-classification settings [11]. Their performance remains context-dependent, and their scores can be difficult to interpret mechanistically [12, 13]. Current evidence therefore supports their use mainly as complementary components of structure-prediction and virtual-screening workflows, rather than as stand-alone methods for affinity prediction or compound prioritisation.

The variable performance of these models reflects not only the difficulty of predicting protein– ligand binding modes and binding affinity, but also the need for large, high-quality datasets for training, fine-tuning, and evaluating structure-based methods. Experimentally determined protein– ligand complexes remain central to model development, while associated affinity measurements are required to connect predicted binding modes to quantitative binding strength. Existing resources such as PDBbind [14] have provided an important foundation, and newer datasets have sought to expand the scale of available protein–ligand data [15–17]. However, many such resources are assembled retrospectively from multiple public sources or achieve scale by augmenting with predicted structures, rather than using experimental data generated through a consistent pipeline. As a result, they can contain substantial noise from predicted data or heterogeneous experimental conditions, assay formats, and curation procedures, complicating both model training and evaluation [18, 19].

These issues highlight the need for novel and experimentally coherent datasets. Dense experimental datasets generated within discovery campaigns provide one route to more robust model training and evaluation. By linking related chemical series with experimentally determined binding modes and affinity measurements, such datasets enable evaluation of model behaviour across local structure–activity relationships, receptor conformational changes, and follow-on compound optimisation [20, 21]. They also provide a practical setting for testing whether target-specific data can improve structure-based models through fine-tuning or other adaptation strategies.

We have therefore established OpenBind as an open-science initiative designed to generate and release coherent experimental structure–affinity datasets for structure-based AI and drug discovery. Our first data release focuses on enteroviral 2A protease, a conserved protease involved in viral polyprotein processing and host-cell modulation [22]. Enterovirus A71 (EV-A71) and the closely related Coxsackievirus A16 (CVA16) are important non-polio enteroviruses that cause hand, foot, and mouth disease [23]. Infection can lead to serious neurological complications in severe cases [24], making 2A protease a relevant antiviral target for structure-guided ligand discovery.

This paper describes the generation, curation, and benchmarking of the first OpenBind structure– affinity dataset for enteroviral 2A protease. It combines crystallographic fragment screening [25], follow-on compound optimisation, and associated affinity measurements (Figure 1). In total, the release contains 925 crystallographic binding events from 699 compounds, together with affinity measurements for 601 compounds. This provides a dense target-focused resource linking experimentally observed binding modes to quantitative binding behaviour across related chemical series.

**Figure 1:**
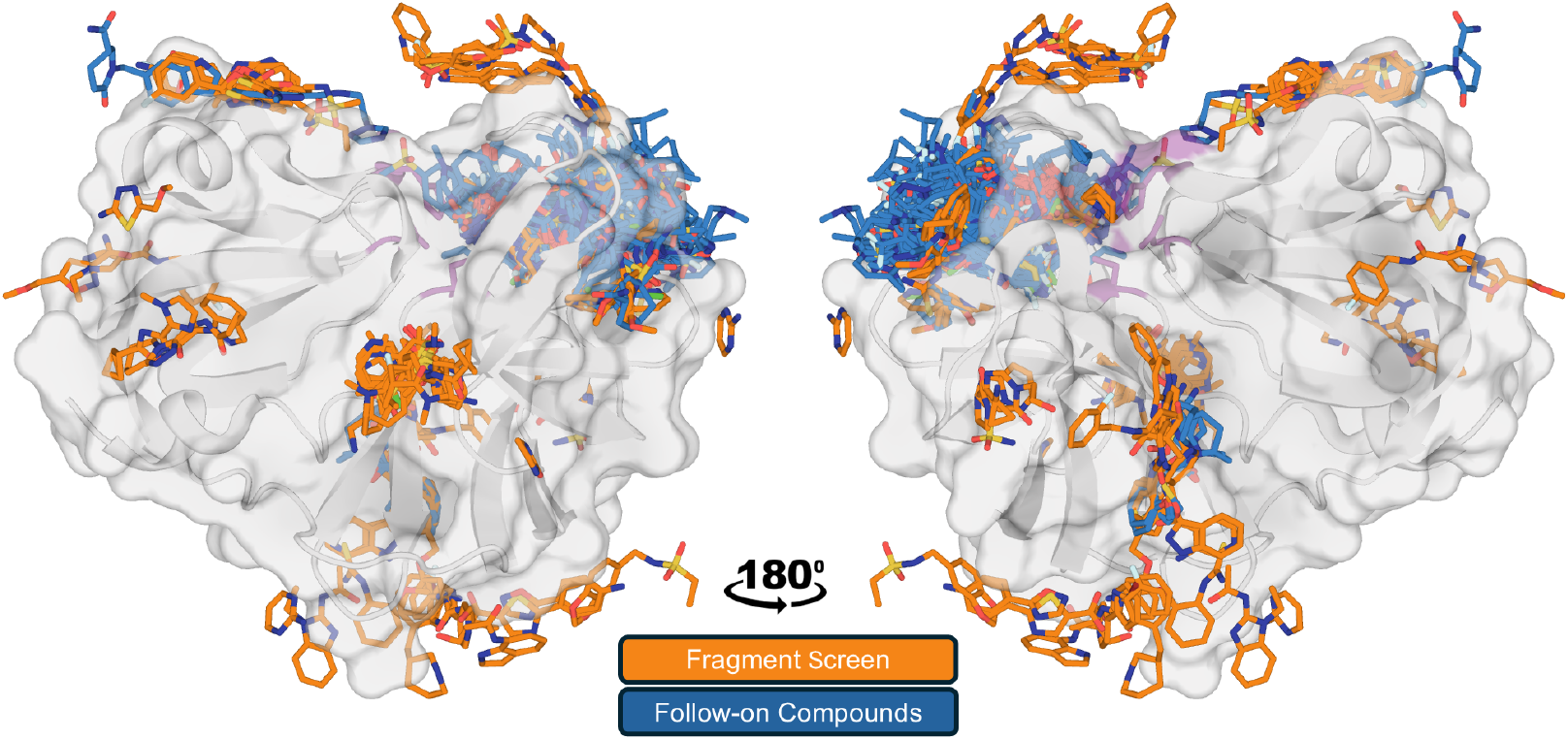
Overview of the OpenBind enteroviral 2A protease release. The protein is shown in surface representation with experimentally observed ligand binding modes overlaid. Fragment-screen binding events (*n* = 113) are shown in orange and follow-on binding events (*n* = 812) in blue. The protease catalytic triad is highlighted in magenta.

To the best of our knowledge, this represents the largest public single-target experimental structure– affinity dataset. OpenBind complements other prospective open data-generation initiatives addressing related but distinct data needs. OpenADMET focuses on experimental data for ADMET-relevant anti-targets [26], whereas Target2035 aims to generate protein–ligand screening data across a diverse set of human proteins, primarily large-scale affinity-selection mass spectrometry measurements and DNA-encoded library screening to identify binders and non-binders [27, 28]. OpenBind instead focuses on dense, target-specific datasets linking experimentally determined structures with quantitative affinity measurements across coherent discovery campaigns.

We use the dataset to assess representative docking, cofolding, virtual screening, fine-tuning, and affinity-prediction approaches in a realistic target-focused setting. Critically, we find that receptor-state selection remains a major determinant of docking performance, that cofolding methods can improve over fixed-receptor cross-docking but still struggle to consistently recover experimentally observed binding modes, and that affinity prediction remains highly challenging across all approaches tested. Notably, fine-tuning on fragment-screen structures substantially improves target-specific cofolding performance, demonstrating that dense experimental datasets can support not only bench-marking and error analysis, but also data-driven model adaptation and improvement.

## 2 Methods

Here, we describe the experimental workflows used to generate the dataset, the curation and benchmark definitions for downstream evaluation, and the docking, cofolding, fine-tuning, and affinity-prediction methods assessed in this work. Additional implementation details, extended parameter settings, and supporting protocols are provided in the Supplementary Information (SI) and accompanying public repositories.

### 2.1 Experimental data collection Target system

The structural and affinity datasets were generated using the 2A cysteine protease from Coxsackievirus A16 (CVA16) strain G10, which served as a surrogate system for Enterovirus A71 (EV-A71). The CVA16 and EV-A71 2A proteases differ by five amino acids, none of which are located within the catalytic site or are expected to substantially affect enzymatic activity [25]. Protein production, purification, and construct-specific details are provided in the SI.

#### Crystallographic screening

The structural dataset was generated from an initial crystallographic fragment screen followed by soaking of follow-on compounds. In total, 1,146 unique fragments and 3,245 unique follow-on compounds were tested by crystallographic soaking. These experiments yielded 113 fragment-derived crystallographic binding events and 812 follow-on binding events. Experimental procedures for crystallisation and crystallographic screening are described in protocols.io [29], with further methodological details provided in the SI.

#### Affinity measurements

Binding affinities were measured using grating-coupled interferometry (GCI) on a Creoptix WAVEsystem employing waveRAPID kinetics [30]. Detailed assay optimisation, protein immobilisation, buffer conditions, and experimental procedures are provided in the SI and associated protocols.io workflow [31].

Affinity measurements were collected for 601 compounds. Sensorgrams were analysed using the standard Creoptix software workflow and manually reviewed by trained experimentalists. Association rates (*k*_*a*_), dissociation rates (*k*_*d*_), equilibrium dissociation constants (*K*_*D*_), and associated fit-quality metrics were derived using a 1:1 binding model. We primarily used a quality-controlled subset of these measurements, as described below in “Affinity benchmark construction”.

### 2.2 Dataset curation and benchmark construction

#### Structural benchmark construction

Ligand-binding events were identified either using PanDDA2 [32] or, for later follow-on datasets in which ligands were clearly visible in conventional 2*F*_*o*_*−F*_*c*_ electron-density maps, by direct inspection and modelling. The resulting protein–ligand complexes underwent additional quality-control checks before downstream computational analysis.

Ligand poses were evaluated using PoseBusters [33], which applies standardised checks for chemical and geometric plausibility, including bond geometry, internal clashes, and protein–ligand steric clashes. Complexes failing at least one PoseBusters check were excluded from the structural benchmark.

Complexes were checked for potential crystal-packing interactions by generating symmetry-related crystal mates in PyMOL [34] and identifying protein atoms within 6 Å of the bound ligand. Complexes identified by this procedure were then manually inspected to assess whether the observed binding pose was likely to represent a crystal-packing artefact.

Following these curation steps, the structural benchmark contained 881 crystallographic binding events: 79 fragment-bound complexes and 802 follow-on complexes. These represented 677 unique compounds in total, comprising 66 fragment-screen compounds and 611 follow-on compounds. Unless otherwise stated, the docking and cofolding benchmarks were performed on the 802 curated follow-on binding events, with the initial fragment-screen hits excluded.

#### Similarity analysis against public structural datasets

To assess the novelty of the OpenBind enteroviral 2A protease dataset relative to existing public protein–ligand structural data, similarity analyses were performed against PLINDER [15]. The primary reference set comprised PLINDER structures deposited on or before 30 September 2021, corresponding to the reported AlphaFold3 training cutoff [5]. Additional similarity distributions using alternative reference cutoffs and dataset subsets are provided in the SI.

Similarity was quantified using the ligand–pocket similarity framework introduced in Runs N’ Poses [6], which combines pocket query coverage with ligand overlap based on molecular shape and pharmacophoric colour. Scores range from 0 to 100, with larger values indicating greater similarity to previously available protein–ligand complexes.

#### Affinity benchmark construction

The release contains Creoptix WAVEsystem [30] affinity measurements for 601 compounds with associated structural data, corresponding to 804 of the 925 crystallographic binding events. To construct the affinity-prediction benchmark, we excluded control measurements and filtered sample measurements using predefined quality-control criteria based on fit boundaries, parameter uncertainty, goodness of fit, and response magnitude, as described in the SI. This procedure yielded affinity measurements for 494 compounds, which was reduced to 490 after excluding binding events removed during the structural benchmark curation.

#### Virtual screening benchmark construction

We constructed a virtual screening benchmark from curated crystallographic binders and suspected non-binders from the soaking campaign. Here, “suspected non-binder” denotes a compound for which no crystallographic binding event was observed in a technically successful soaking experiment, rather than a biochemical determination of binding status. Compound structures were standardised, deduplicated at the compound level, and filtered using the experimental quality-control and subset-definition criteria detailed in the SI.

The benchmark was restricted to follow-on compounds, excluding compounds from the fragment-screen collections and small fragment-like compounds. This resulted in *n* = 2,084 compounds, comprising 566 binders and 1,518 suspected non-binders. Potential physicochemical and chemical-space differences between binders and suspected non-binders were examined using selected RD-Kit descriptor distributions, nearest-neighbour comparisons to both binding and non-binding compounds, and UMAP dimensionality reduction of ECFP4 fingerprints.

### 2.3 Computational benchmarking and analysis Structural benchmark metrics

Physical plausibility of predicted protein–ligand complexes was assessed using PoseBusters v0.6.5 [33]. Poses passing all PoseBusters checks were classified as PoseBusters-valid (PB-valid). Pose-Busters validity was treated as a prerequisite for all reported success metrics.

Pose accuracy and protein–ligand interaction recovery were evaluated with OpenStructure v2.11.1 [35] using binding-site-superposed, symmetry-corrected heavy-atom ligand RMSD and LDDT-PLI scores, respectively. Two structural success criteria were reported throughout: an RMSD-based success criterion required a PB-valid complex and ligand RMSD *≤* 2.0 Å (“RMSD-valid”), while a stricter joint success criterion required a PB-valid complex, ligand RMSD *≤* 2.0 Å, and LDDT-PLI *≥* 0.8 (“Success”).

Performance was evaluated at different rank cutoffs, primarily Top-25 and, where indicated, Top-1. Top-1 performance assesses whether the highest-ranked prediction satisfies the success criterion. Top-25 performance was considered successful when at least one of the 25 retained predictions satisfied the corresponding success criterion, and therefore evaluates whether a method can generate a correct pose even if it does not rank it first.

#### Docking benchmarks

Docking benchmarks were performed using GNINA v1.3.2 [36], smina [37] with the Vina v1.1.2 scoring function [38], and DiffDock-L v1.1.3 [39]. Each method was evaluated in four receptor scenarios: redocking into the cognate crystallographic receptor, cross-docking into two apo-like receptor structures, and fragment-informed cross-docking using the receptor from the most chemically similar fragment-screen complex.

For apo cross-docking, an experimentally determined apo structure (PDB ID: 8POA) and an AlphaFold2-predicted apo model obtained from the AlphaFold Protein Structure Database (AFDB entry AF-0000000365760404) [40], associated with the Viro3D viral protein structure resource [41], were used. For fragment-informed cross-docking, each follow-on compound was assigned the receptor from the fragment-screen complex with the highest ECFP4 Tanimoto similarity [42].

GNINA and smina generated poses from five prepared ligand conformers, which were pooled and ranked using the corresponding docking score. DiffDock was run five times per complex, and predictions were ranked by confidence score. For each method and receptor scenario, the 25 highest-ranked poses were retained and evaluated using the structural benchmark metrics described above. Full ligand and receptor preparation procedures and docking parameters are provided in the SI.

#### Cofolding benchmarks

Cofolding benchmarks were performed using AlphaFold3 [5], Boltz-1 [43], Boltz-2 [8], OpenFold3-p2 [44], Protenix-v1 [45], and RosettaFold3 [46]. Each method received the target protein sequence, ligand SMILES string, and a CCD code specifying a single zinc ion as input. The protein was treated as a monomer in all cofolding predictions.

To reduce differences arising from multiple sequence alignment (MSA) generation, all models were run using the same precomputed MSA, generated from the ColabFold database using the OpenFold3 MSA generation pipeline. All evaluated cofolding models, except Boltz-2, were trained on structural data up to the AlphaFold3 training-data cutoff of 30 September 2021. Boltz-2 was evaluated using the available public weights, which were trained on structures released up to 1 June 2023.

For each protein–ligand complex, five independent predictions were generated using different random seeds. Each prediction produced five candidate structures, yielding 25 predicted complexes per target. Predicted complexes were ranked using the method-specific confidence score associated with the protein–ligand interface and evaluated using the benchmark metrics described above.

#### OpenFold3 fine-tuning

To investigate the impact of fragment-derived structural information on follow-on compound prediction, we fine-tuned OpenFold3-p2 on the curated set of fragment-bound structures (*n* = 79) and evaluated on the follow-on compound benchmark using the same protocol as the base model. Fragment-bound structures that had been solved with two copies in the asymmetric unit were processed so that only the fragment-bound copy was retained. The model was fine-tuned using a learning rate of 1*×*10^*−*3^ (100-step linear warm-up) and a batch size of 8 for 1,000 total training steps, requiring approximately 8 h of wall-clock time on 8 NVIDIA GH200 GPUs.

#### Affinity prediction benchmark

Affinity predictions were evaluated using AEV-PLIG [47], GNINA [36], smina [37], AQAffinity [9], and Boltz-2 [8]. For AEV-PLIG, affinities were predicted from the experimentally determined protein–ligand complexes. For GNINA and smina, predicted affinities were derived from the highest-scoring redocked pose. For AQAffinity and Boltz-2, the protein–ligand complex was predicted by the model, and the corresponding predicted affinity was used for evaluation.

All model outputs were converted to pK units, with larger values indicating stronger predicted binding, to enable comparison with the experimental pK_D_ values. Evaluation was performed at the compound level. When multiple structures or predictions were available for the same compound, the corresponding predicted affinities were averaged before calculating the evaluation metrics.

#### Virtual screening benchmark

Virtual screening performance was evaluated on the full benchmark set (*n* = 2,084), comprising 566 binders and 1,518 suspected non-binders. For docking-based methods, receptor selection followed the fragment-informed protocol used in the structural benchmark. GNINA and smina docking followed the structural benchmark protocol, except compounds were ranked by GNINA using CNN VS rather than CNNscore, as CNN VS has been reported to be better suited to compound ranking in virtual-screening settings [48]. For cofolding-based virtual screening, the base and fragment-fine-tuned OpenFold3-p2 models were evaluated using the same prediction protocol as in the structural benchmark, with compounds ranked by pair ipTM scores.

Performance was assessed primarily using the area under the receiver operating characteristic curve (ROC-AUC), with average precision and enrichment factors at 1%, 2%, and 5% reported in the SI.

## 3 Results and Discussion

The first OpenBind enteroviral 2A protease release provides a dense experimental structure–affinity dataset for evaluating structure-based modelling methods on a coherent antiviral discovery campaign. We examined several related questions: how novel the complexes are relative to existing public structural datasets (Section 3.1), how docking performance depends on receptor-state selection (Section 3.2), whether cofolding methods can accurately accommodate receptor conformational flexibility (Section 3.3), whether fragment-screen structures can improve target-specific modelling through fine-tuning (Section 3.4), how well current approaches relate structural information to measured binding affinity (Section 3.5), and the ability of methods to identify binders (Section 3.6).

Unless otherwise stated, benchmark analyses were performed on the curated follow-on compound set, comprising 802 protein–ligand complexes that passed PoseBusters checks and were not flagged for potential crystal-packing effects.

### 3.1 Similarity to existing structural datasets

A central question for any new structural data release is whether it provides information beyond that already available in public protein–ligand datasets. This is important for both model training and evaluation, since strong similarity to structures available during model development can make it difficult to distinguish genuine generalisation from performance driven by near-neighbour information. Runs N’ Poses demonstrated this directly, showing that cofolding performance decreases substantially as similarity to previously available structures declines [6].

To assess the novelty of the OpenBind enteroviral 2A protease release, we compared the curated complexes against public protein–ligand structures released on or before 30 September 2021, corresponding to the reported structural training cutoff for AlphaFold3 and several related cofolding models. Similarity was quantified using the Runs N’ Poses ligand–pocket metric, which combines pocket coverage with ligand shape and pharmacophoric-feature overlap [6]. Scores range from 0 to 100, with higher values indicating greater similarity to previously available complexes. In the Runs N’ Poses benchmark, cofolding success was typically approximately 10–40% below a score of 50, compared with approximately 80–90% for structures with a similarity score between 80 and 100.

The OpenBind complexes showed only moderate similarity to structures available before this cutoff (Figure 2). Across all 881 curated crystallographic binding events, the median score was approximately 40, and almost all complexes scored below 50. This places the dataset in a similarity regime where Runs N’ Poses observed substantial declines in cofolding performance, making it a useful test of model generalisation beyond closely related training examples. Fragment-bound structures showed a broader distribution and generally lower similarity scores. This could reflect their limited representation in public protein–ligand datasets, limitations of the measure in scoring fragments contained within larger bound compounds, or both. Full summary statistics and analyses using alternative structural cutoffs are provided in the SI (Figure S1; Table S1). Together, these results indicate that the OpenBind complexes were not strongly represented in public structural data available before the reference cutoff.

**Figure 2:**
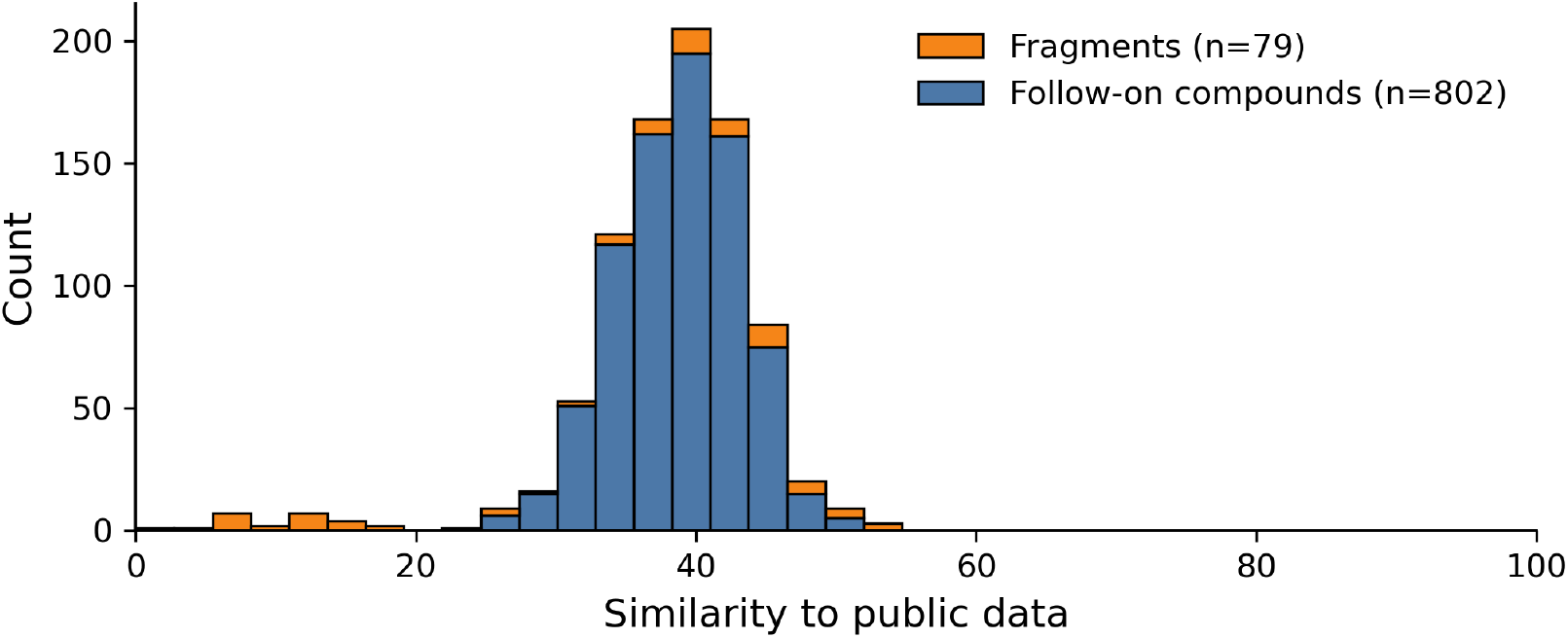
Distribution of ligand–pocket similarity scores between curated OpenBind enteroviral 2A protease complexes and public protein–ligand structures released on or before 30 September 2021. Fragment-bound and follow-on complexes are shown in orange and blue, respectively. Similarity was measured using the ligand–pocket similarity metric from Runs N’ Poses [6].

We also examined the chemical similarity between the fragment hits and follow-on compounds. For each of the 611 unique curated follow-on compounds, the ECFP4 Tanimoto similarity to every curated fragment was calculated, and the compound was assigned to the fragment with the highest similarity. These nearest-fragment assignments were concentrated among a small number of fragments: x0926a was the most similar fragment for 507 compounds (83.0%), while x0812a and x1019a were the closest fragments for a further 30 (4.9%) and 14 compounds (2.3%), respectively. Several examples are shown in Figure 3 and complete assignment counts are provided in Figure S2.

**Figure 3:**
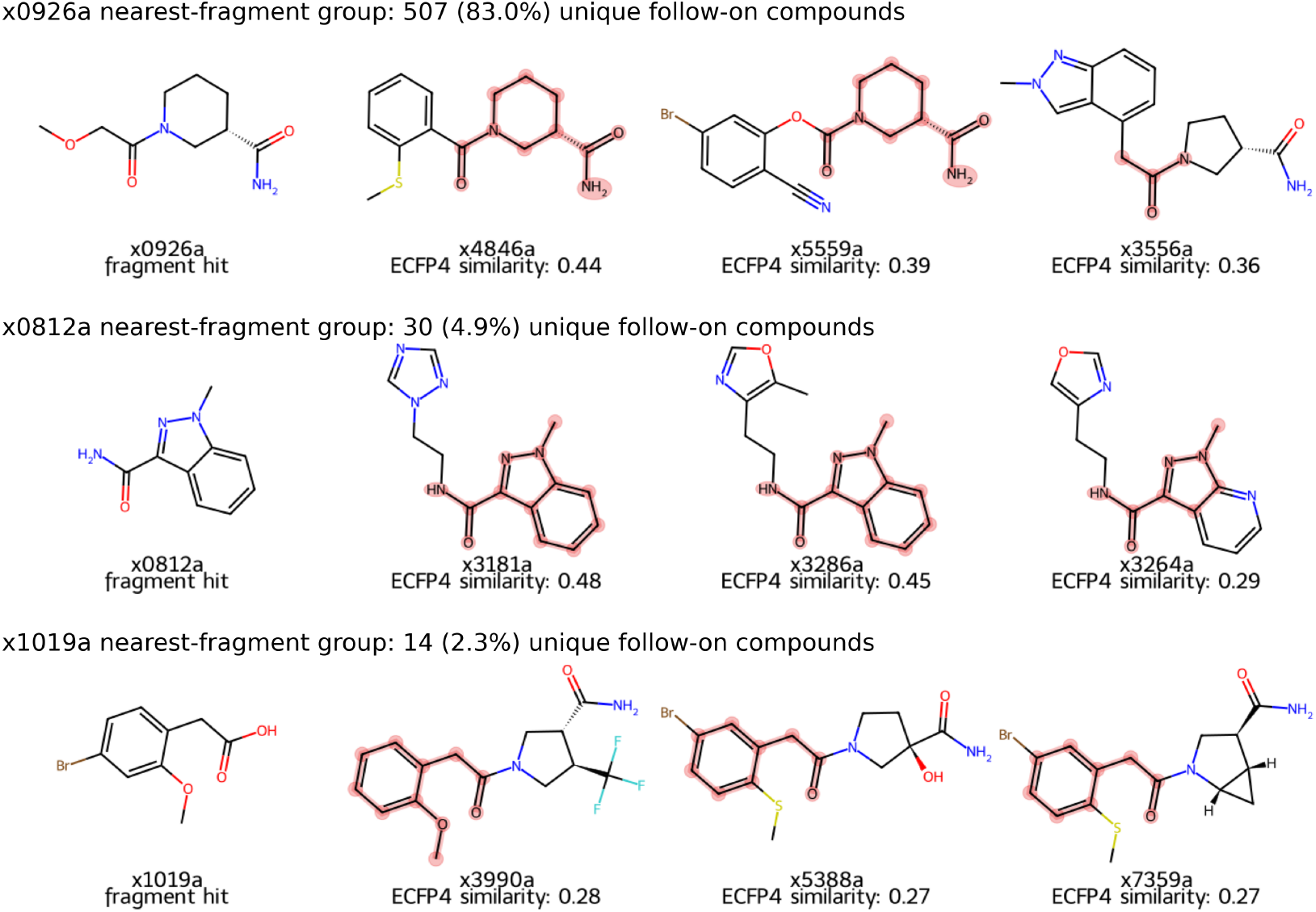
Representative fragment hits and follow-on compounds from the three largest nearest-fragment groups. In each group, the leftmost structure is the fragment hit, followed by representative follow-on compounds selected across the corresponding ECFP4 Tanimoto similarity distribution. Highlighted atoms and bonds indicate the maximum common substructure shared between each follow-on compound and the fragment hit.

Despite this concentration, the maximum ECFP4 Tanimoto similarities within the three largest nearest-fragment groups showed substantial variation, spanning a range of approximately 0.2–0.6, with a median just below 0.4 (Figure S3). Representative compounds from these groups further show substantial chemical variation beyond the shared substructures, including differences in molecular size, substitution pattern, ring systems, and functional-group composition (Figure 3). The dataset therefore contains chemically coherent nearest-fragment groups while retaining considerable variation within them.

These analyses should not be interpreted as complete measures of benchmark difficulty or medicinal-chemistry provenance. Similarity to public structures is only one determinant of model performance, and the ligand–pocket similarity metric used here represents one approach to estimating similarity to previously available public structural data. Other factors, such as ligand chemistry, receptor flexibility, and evaluation design, also contribute to model performance. Likewise, nearest-fragment assignments summarise chemical relatedness but do not necessarily reflect formal synthetic lineage. Nevertheless, the results show that the release combines limited similarity to earlier public structural data with internally coherent fragment-related chemical series.

### 3.2 Docking benchmarks

Docking performance depended strongly on the receptor conformation used for prediction (Figure 4). In the redocking setting, where the experimentally observed receptor conformation was provided, the best-performing methods achieved strong pose recovery rates across the benchmark set. GNINA achieved a Top-25 success rate of approximately 85% across the follow-on compounds, indicating that current docking approaches can often recover experimentally observed binding modes when the receptor already adopts a ligand-compatible conformation.

**Figure 4:**
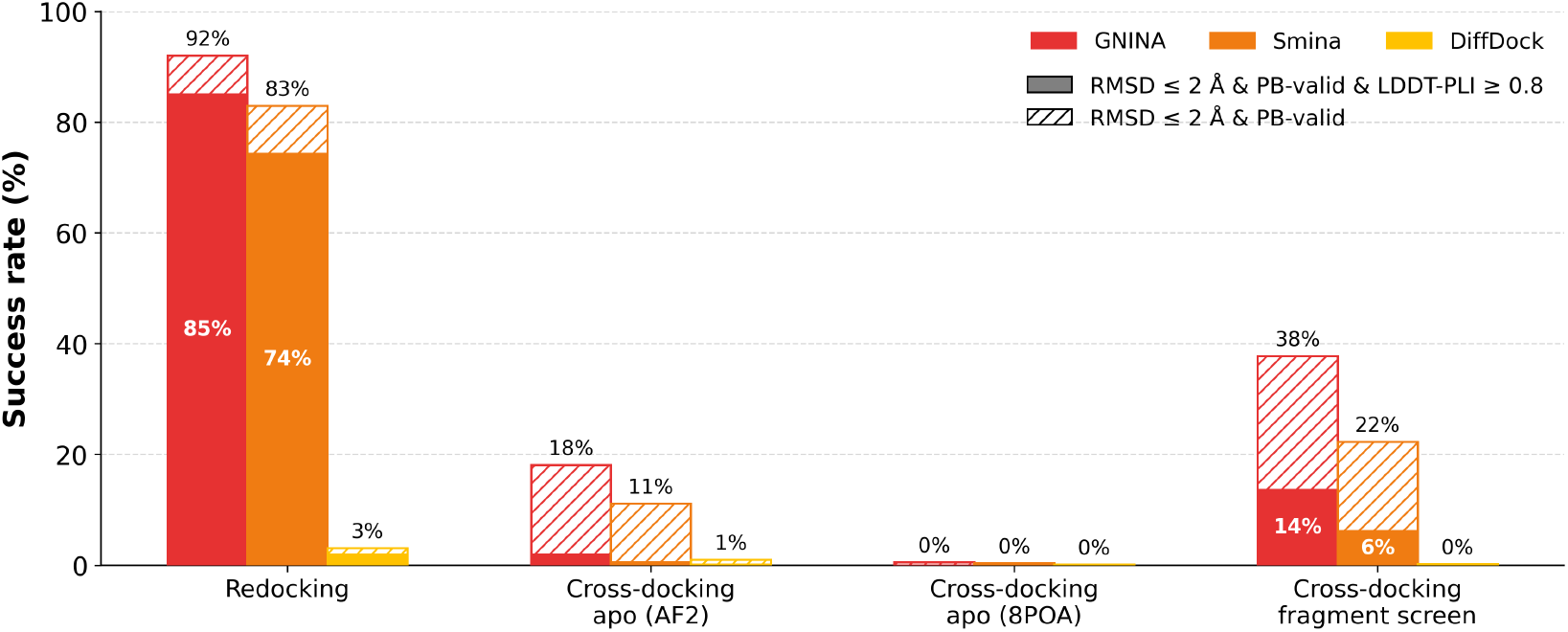
Top-25 docking performance on the curated follow-on benchmark (*n* = 802 binding events). The solid portion of each bar shows the fraction of predictions that recovered the experimental binding mode with ligand heavy-atom RMSD *≤* 2 Å, LDDT-PLI *≥* 0.8, and PoseBusters validity. Hatched portions show the additional predictions that met the RMSD and PoseBusters criteria but not the LDDT-PLI threshold. Docking performance is substantially stronger when cognate receptor conformations are available (redocking), while cross-docking into apo structures is considerably more challenging. Using receptor conformations derived from chemically related fragment-bound structures partially improves pose recovery but remains substantially below redocking performance, highlighting the importance of receptor-state selection.

Cross-docking proved substantially more challenging. We docked ligands into an experimentally determined apo structure (PDB ID: 8POA) and the predicted apo structure from the AlphaFold Protein Structure Database [40, 41]. In both cases, success rates dropped substantially for all evaluated methods. Structural comparison of ligand-bound and apo conformations revealed substantial rearrangement of a loop adjacent to the binding site (Figure 5). In the apo conformations, this loop partially occludes the ligand-binding pocket and prevents recovery of many crystallographic binding modes without substantial steric clashes. The poor cross-docking performance therefore appears to arise primarily from receptor-state mismatch rather than failures in ligand placement or scoring alone.

**Figure 5:**
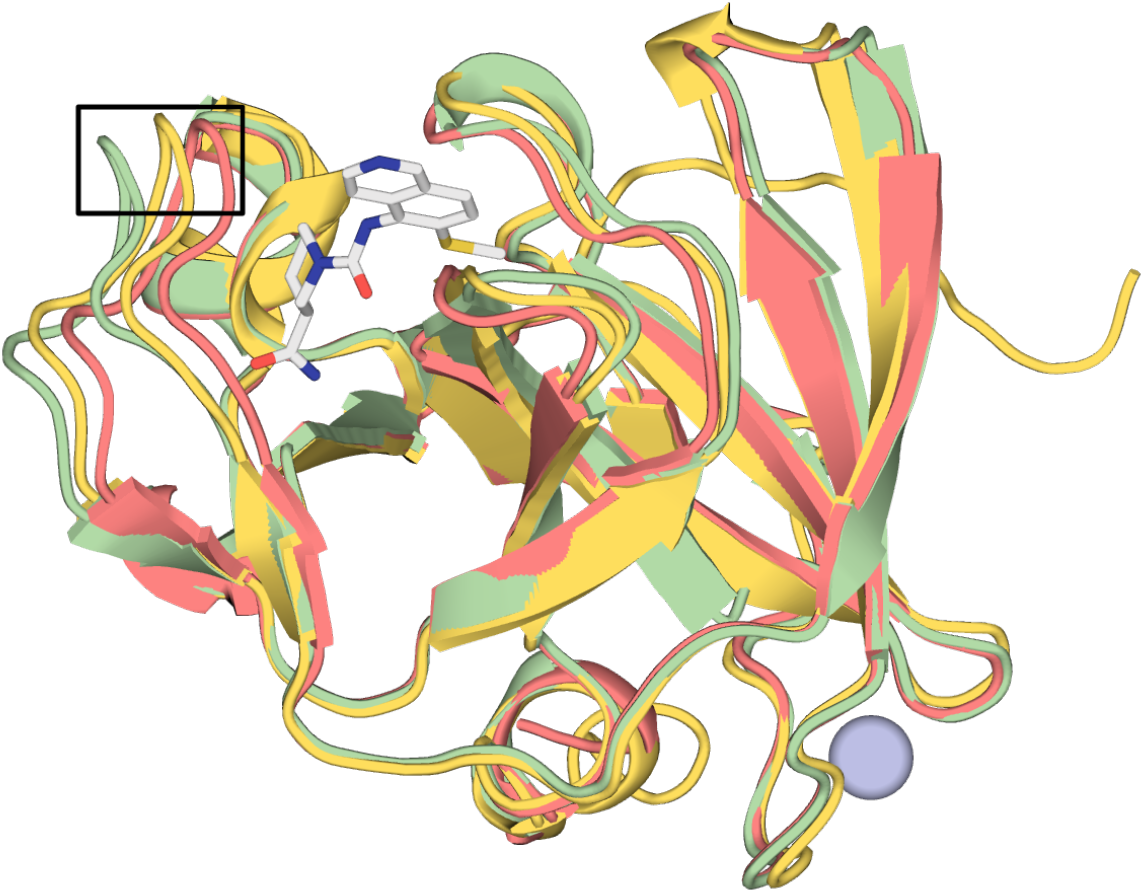
Comparison of ligand-bound and apo conformations of enteroviral 2A protease. The ligand-bound structure (Fragalysis code x5261a, green), experimentally determined apo structure (PDB ID 8POA, red), and AlphaFold2-predicted apo structure (yellow) adopt distinct conformations in a loop adjacent to the ligand-binding pocket, highlighted by the black box. The ligand is shown in white, and the Zn^2+^ ion as a light-blue sphere. In both apo structures, this loop partially occludes the binding site, contributing to substantially reduced rigid-body cross-docking performance.

Fragment-bound receptor structures are more likely than apo-like ones to capture ligand-compatible pocket geometries, and chemically related fragments may occupy similar subpockets or induce local loop and side-chain arrangements that better accommodate related ligands. For enteroviral 2A proteases, this is particularly relevant because the binding pocket is partially closed in the apo structures, making rigid docking difficult or sterically incompatible for many ligands. Fragment-informed cross-docking improved GNINA RMSD-valid performance to 38%, indicating that a more compatible receptor state often enables approximate pose recovery. However, only 14% of compounds also satisfied the stricter success criterion, which requires LDDT-PLI *≥* 0.8, suggesting that the detailed protein–ligand contact pattern is not consistently recovered. This may reflect the smaller size of the fragment ligands, which can capture only part of the pocket conformation required by larger follow-on compounds. Docking against multiple fragment-informed receptor structures substantially improved pose recovery (Top-25 RMSD-valid 54%, Success 23% for five fragment receptors; Figure S4), although this comes at additional computational cost and the performance improvements appear to diminish as more structures are added.

Together, these results highlight receptor-state selection as a major determinant of docking performance for the enteroviral 2A protease system. The dataset therefore provides a realistic benchmark for evaluating methods designed to model binding-site flexibility and local loop rearrangements in practical structure-based drug discovery workflows.

### 3.3 Cofolding benchmarks

The cross-docking results highlight a major challenge in structure-based modelling: if the receptor adopts an incompatible binding-site conformation, recovering the experimentally observed ligand pose may be impossible for rigid-body docking. Cofolding methods offer an alternative approach to this problem by jointly predicting receptor and ligand conformations, potentially allowing the binding site to adapt during complex prediction.

We also evaluated the performance of several recent cofolding methods, namely AlphaFold3, Boltz-1, Boltz-2, RosettaFold3, Protenix-v1, and OpenFold3-p2. Predictions were evaluated using the same criteria described for the docking benchmarks.

Cofolding substantially improved success rates relative to rigid cross-docking into apo receptor conformations (Figure 6). Unlike rigid cross-docking, cofolding is not constrained by the partially closed apo-like receptor structures, but can instead predict a more ligand-compatible pocket geometry. Several cofolding methods generated relatively accurate binding-pocket conformations, although this did not always translate into correct ligand placement (Figure S5).

**Figure 6:**
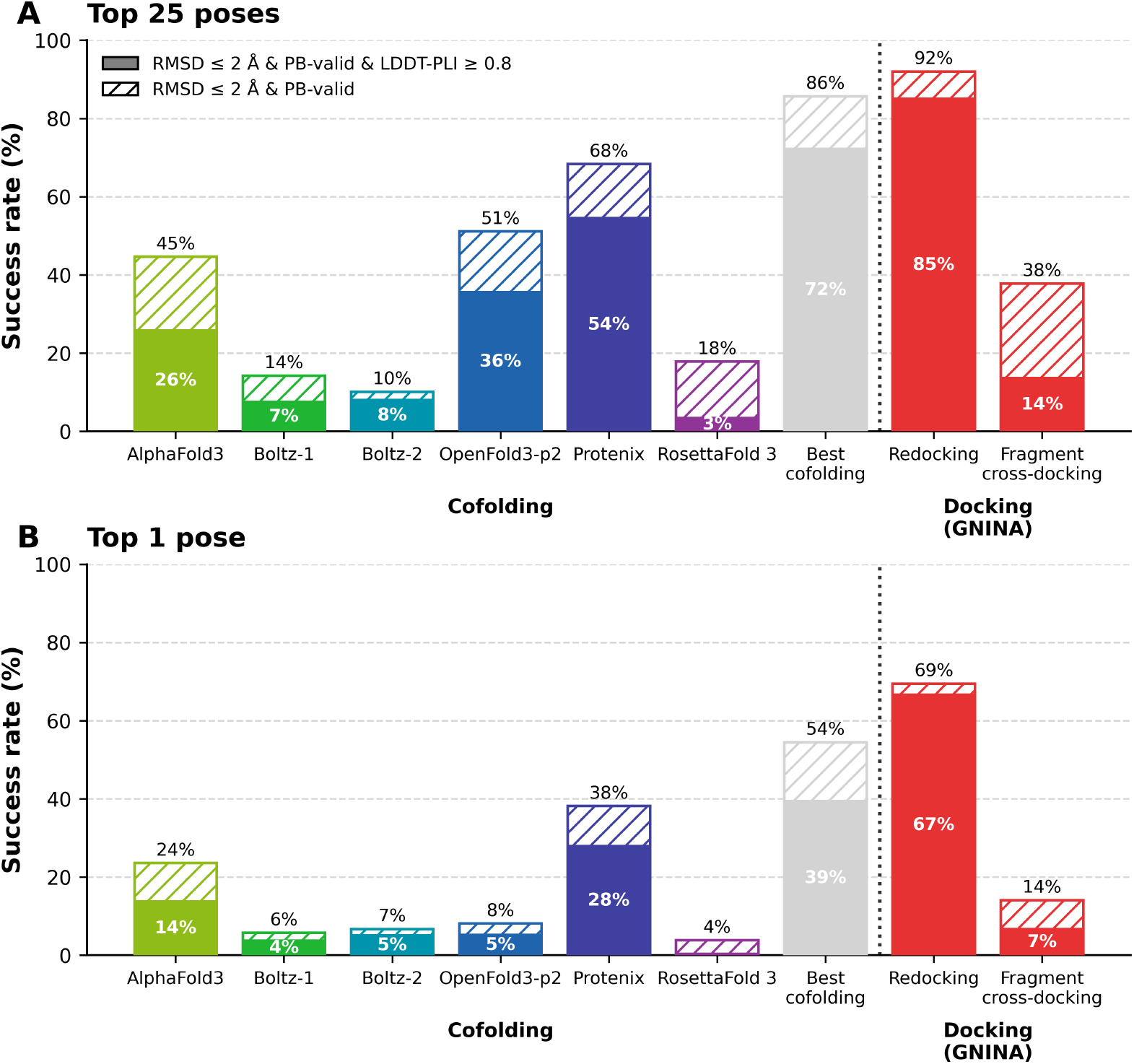
Performance of representative cofolding methods on the curated follow-on benchmark (*n* = 802 binding events), shown for the Top-25 (A) and Top-1 (B) settings. Solid bars indicate predictions satisfying heavy-atom RMSD *≤* 2 Å, LDDT-PLI *≥* 0.8, and PoseBusters validity, while hatched extensions indicate additional predictions satisfying RMSD *≤* 2 Å and PoseBusters validity only. “Best cofolding” denotes an oracle across the top 25 poses from all tested cofolding methods (A), and across the top-ranked pose from each method (B). GNINA docking into the cognate holo structure (redocking) and fragment-informed cross-docking are shown for reference.

Among the individual cofolding methods, Protenix-v1 achieved the strongest Top-25 performance, followed by OpenFold3-p2 and AlphaFold3, while Boltz-1, Boltz-2, and RosettaFold3 showed lower overall success rates. Performance decreased substantially for all cofolding methods when only the top-ranked prediction was considered, with Protenix-v1 retaining the strongest Top-1 performance, followed by AlphaFold3. This difference between Top-25 and Top-1 results indicates that generating an accurate complex and ranking it as the highest-confidence prediction remain distinct challenges. As these results are derived from a single protein target, the relative ordering of methods and the balance between docking and cofolding may not generalise to other systems.

GNINA redocking and fragment-informed cross-docking are included for reference. Redocking does not represent a prospective prediction scenario, as it uses the cognate ligand-bound receptor conformation, but provides a strong reference for pose recovery when the appropriate receptor state is known. Fragment-informed cross-docking instead uses a single experimentally determined receptor from the initial fragment screen selected by ligand similarity, and thus more closely resembles prospective use. Both docking approaches retained a larger proportion of their Top-25 performance at the Top-1 level than most cofolding methods.

An oracle “best-of” cofolding ensemble achieved substantially higher success rates than any individual method, indicating that different methods succeed on different subsets of complexes. However, this upper bound was obtained across a substantially larger number of generated predictions and therefore does not directly correspond to a practical inference setting. Nonetheless, the result suggests that improved ranking, confidence estimation, or ensemble-selection strategies may substantially improve practical cofolding performance. Failure-mode analysis further indicates that accurate ligand pose generation remains challenging for most methods, even when the protein pocket is identified and modelled correctly (Figure S5).

Overall, these results indicate that cofolding methods can partially overcome the receptor conformational mismatch that limits rigid cross-docking approaches, but that accurately recovering and ranking experimentally observed receptor–ligand interaction geometries remains challenging for many complexes within this benchmark (Figures 6, S5). These findings are broadly consistent with recent benchmarking studies on newly determined protein–ligand complexes, which reported that modern cofolding methods can often recover ligand poses but still face challenges in ranking, affinity estimation, and virtual screening performance [11]. The OpenBind enteroviral 2A protease dataset therefore provides a useful prospective test case for evaluating how effectively current cofolding approaches generalise to experimentally novel protein–ligand systems and chemically diverse follow-on compounds.

### 3.4 OpenFold3 fine-tuning

The structure and organisation of the OpenBind dataset additionally enable investigation of whether fragment-derived structural information can improve prediction performance on related follow-on compounds. Fragment-based drug discovery can efficiently sample chemical space and provide experimentally resolved binding modes that guide the optimisation of larger, higher-affinity compounds [49]. To explore whether these advantages can also be leveraged by structure-prediction models, we fine-tuned OpenFold3-p2 using fragment-bound structures from the OpenBind release and evaluated its performance on the follow-on compound benchmark.

Fine-tuning on fragment-derived structural data substantially improved performance on the follow-on compound benchmark relative to the base OpenFold3-p2 model (Figure 7). The largest improvements were observed for the stricter combined success criteria incorporating both RMSD and LDDT-PLI, indicating that fine-tuning improved not only approximate ligand placement but also recovery of detailed protein–ligand interaction geometry. These results suggest that experimentally determined fragment-screen structures can provide a useful target-specific learning signal for down-stream protein–ligand modelling tasks.

**Figure 7:**
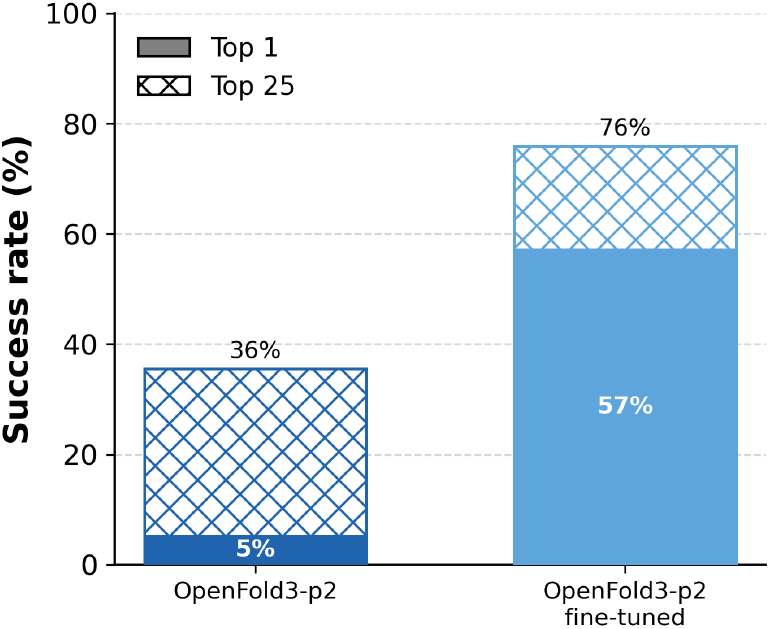
Effect of fragment-derived fine-tuning on OpenFold3-p2 performance for follow-on compounds. Success required ligand heavy-atom RMSD *≤* 2 Å, LDDT-PLI *≥* 0.8, and PoseBusters validity. Solid bars represent Top-1 success rates, while crosshatched bars show Top-25 success rates. Fine-tuning on experimentally determined fragment-bound structures substantially improved performance, suggesting that target-specific structural information from fragment screening can provide a useful signal for downstream modelling tasks.

Because the fine-tuning and evaluation data arise from a single target-focused discovery campaign, in which many follow-on compounds were derived from fragment-screen hits (Figure S2), the extent to which these gains reflect broader transfer across chemical series or protein targets remains to be established.

Nevertheless, for this dataset, the results demonstrate that fragment-derived structural information can be used to improve prediction of related follow-on compounds and illustrate how dense experimental datasets linking fragment screening and follow-on optimisation can support systematic investigation of adaptation and target-specific fine-tuning in structure-based AI models.

### 3.5 Affinity prediction benchmarks

The OpenBind enteroviral 2A protease release additionally contains experimentally measured binding affinities for 601 compounds, corresponding to 804 of the 925 crystallographic binding events. Following quality-control filtering of the biophysical measurements and removal of binding events that failed PoseBusters validation or were identified as suspected artefacts, 490 compounds were retained for the final affinity-prediction analysis.

We evaluated several affinity prediction approaches for their ability to predict and rank binding strength across the enteroviral 2A protease series. These included structure-based methods such as Boltz-2 [8], GNINA [36], and AEV-PLIG [47]. A constant mean p*K*_*D*_ predictor was included as an RMSE baseline, while molecular weight and cLogP were used as simple physicochemical baselines for rank correlation.

Affinity prediction remained challenging across the benchmark dataset. Several methods produced only modest correlations with experimentally measured affinities (Figure 8). Boltz-2 and GNINA achieved the strongest rank correlations among the evaluated structure-based methods, while smina, AEV-PLIG and Boltz-2 produced the lowest RMSE values. Additional analysis of GNINA and smina across crystal rescoring, minimised crystal structures, redocking, and fragment cross-docking is provided in Figure S6. However, a simple molecular-weight baseline achieved the highest overall Spearman correlation and the mean pK_D_ predictor the lowest RMSE.

**Figure 8:**
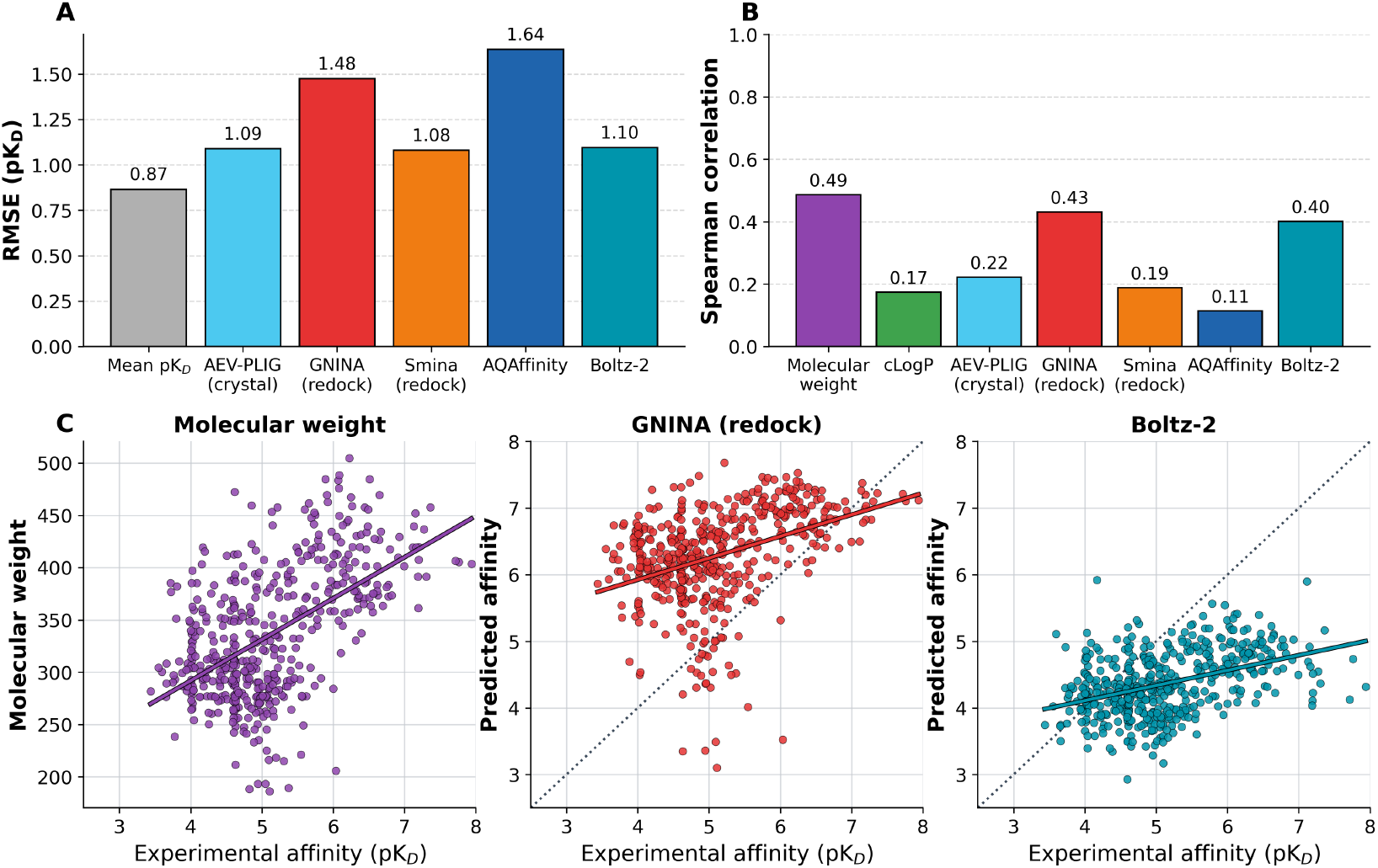
Affinity prediction performance across the OpenBind enteroviral 2A protease benchmark (*n* = 490). **(A)** Root-mean-square error (RMSE) for the evaluated affinity-prediction methods, together with a mean-pK_*D*_ baseline. **(B)** Spearman rank correlation for the evaluated methods and simple physicochemical baselines based on molecular weight and cLogP. **(C)** Relationships between experimentally measured affinity and molecular weight, GNINA redocking predictions, and Boltz-2 predictions. Boltz-2 and GNINA show the strongest rank correlations among the evaluated structure-based methods, whereas smina, AEV-PLIG and Boltz-2 achieve the lowest RMSE values. The strong molecular-weight correlation and the competitive performance of other simple physicochemical baselines highlight the difficulty of extracting accurate affinity predictions from protein–ligand structural information alone.

This result should not be interpreted as evidence that molecular weight alone provides a sufficient model of binding affinity. Rather, it highlights the importance of including simple physicochemical baselines when evaluating affinity-prediction methods, particularly within coherent medicinal chemistry campaigns where potency may correlate with compound elaboration, molecular size, or scaffold progression. Structure-based methods that explicitly model protein–ligand interactions should ultimately be expected to outperform such baselines if they are successfully capturing the structural determinants of affinity.

More broadly, these results illustrate the difficulty of connecting experimentally observed protein– ligand structures to quantitative binding affinity. This OpenBind dataset therefore provides a realistic benchmark for evaluating whether affinity-prediction models can move beyond coarse chemical trends to capture local structure–activity relationships, distinguish productive from non-productive elaborations, and accurately relate binding geometry to experimentally measured potency.

### 3.6 Virtual screening benchmark

To support evaluation of virtual screening methods, we constructed a benchmark from experimentally observed binders and suspected non-binders among the follow-on compounds tested during the soaking campaign. After structure standardisation, deduplication, and filtering, the final benchmark contained 2,084 compounds, comprising 566 binders and 1,518 suspected non-binders.

The resulting benchmark reflects a target-focused discovery campaign rather than a conventional screening collection assembled from chemically distinct active and inactive compounds. Many compounds belong to related chemical series, within which only a subset were experimentally observed to bind. Consistent with this design, an ECFP4-based UMAP embedding showed substantial overlap between binders and suspected non-binders across the sampled chemical space (Figure S7). Regions enriched in binders generally also contained structurally related suspected non-binders, indicating that successful methods must distinguish activity within local chemical series rather than separate globally distinct molecular classes. At the same time, several more isolated regions were populated predominantly by suspected non-binders and were comparatively dissimilar from the binder set, reflecting the broader chemical diversity of the suspected non-binder collection.

The two classes also showed broadly overlapping distributions of basic molecular properties such as molecular weight, heavy-atom count, and hydrogen-bonding capacity (Figure S10). Their close structural relationship was further supported by nearest-neighbour analysis (Figure S8). For each binder, the median ECFP4 Tanimoto similarity to the nearest suspected non-binder was 0.661, and 91.0% of binders had a suspected non-binder neighbour with similarity of at least 0.5. Conversely, the median similarity from each suspected non-binder to its nearest binder was 0.425, with 40.4% having a binder neighbour with a similarity of at least 0.5. Thus, most binders have a close structural analogue among the suspected non-binders, whereas the larger suspected non-binder set additionally covers regions with fewer known binders. Representative nearest-neighbour pairs are shown in Figure S9.

These analyses indicate that the benchmark is not readily separable using coarse physicochemical properties or global fingerprint similarity. Instead, it tests whether a method can rank closely related compounds that differ in experimentally observed binding behaviour.

Virtual screening performance was evaluated on the full benchmark consisting of 2,084 compounds. Full performance statistics, including ROC-AUC, average precision, and enrichment factors at 1%, 2%, and 5%, are reported in Table S2.

The molecular weight baseline provided only weak discrimination between binders and suspected non-binders, with a ROC-AUC of 0.539 (Figure 9). This result is consistent with the overlapping descriptor distributions, although some early enrichment was observed among the highest-ranked compounds.

**Figure 9:**
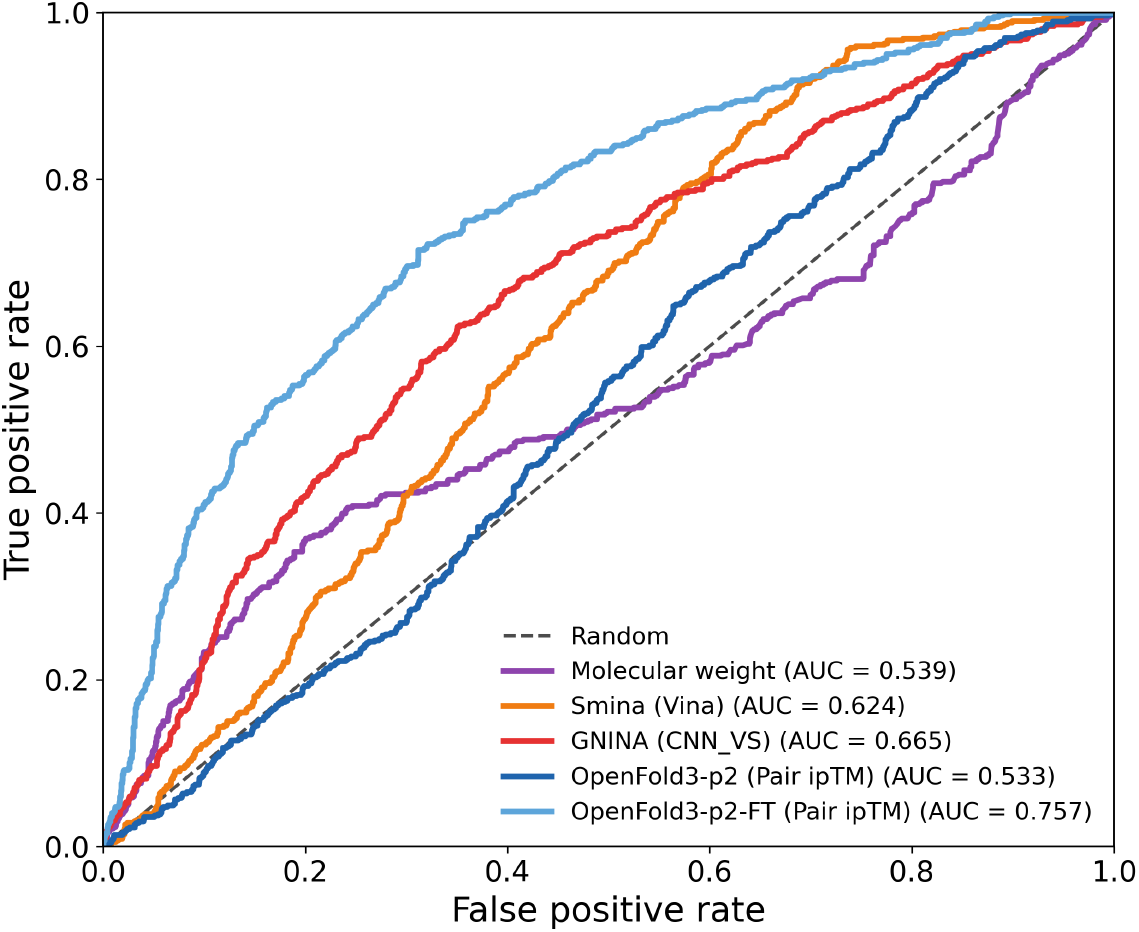
Virtual screening performance on the OpenBind enteroviral 2A protease benchmark (*n* = 2,084; 566 binders and 1,518 suspected non-binders). ROC curves are shown for a molecular-weight baseline, smina, GNINA using CNN VS, and the base and fragment-fine-tuned OpenFold3-p2 models using pair ipTM for both pose selection and compound ranking.

Docking-based methods performed better, although their discrimination remained moderate. Smina, using the Vina scoring function, achieved a ROC-AUC of 0.624. GNINA improved the ROC-AUC to 0.665 when compounds were ranked using CNN VS. The difference between the two methods indicates that the learned GNINA score captures information relevant to binder ranking that is not represented by the conventional empirical scoring function alone.

The base OpenFold3-p2 model performed poorly when pair ipTM was used for compound ranking, achieving a ROC-AUC of 0.53. This is close to the molecular-weight baseline and indicates that, without target-specific adaptation, model confidence in the predicted protein–ligand interface is not a useful proxy for binding in this benchmark. In contrast, the fragment-fine-tuned OpenFold3-p2 model achieved a ROC-AUC of 0.76, exceeding the docking-based methods used in this benchmark.

Because the same pair ipTM-based ranking procedure was applied to the base and fine-tuned models, the observed performance difference is associated with changes introduced by fine-tuning rather than with a change in the evaluation procedure. This suggests that training on target-specific fragment-bound structures made the model’s confidence estimates more informative for distinguishing binders from suspected non-binders in this benchmark. However, pair ipTM remains a confidence metric rather than an explicitly trained affinity or virtual screening score, and its performance should therefore be interpreted as suggesting that it may contain useful ranking signal rather than as evidence of direct binding-affinity prediction. Moreover, because this analysis considers a single target and a single follow-on compound collection, further evaluation across additional targets and chemical series will be required to determine how broadly this improvement generalises.

Overall, conventional docking scores provided moderate discrimination, whereas the unadapted OpenFold3-p2 confidence score contained little useful ranking signal for this dataset. Target-specific fragment fine-tuning substantially improved performance, suggesting that structural information derived from experimentally observed fragment complexes can support the prioritisation of related follow-on compounds. Whether this effect extends to other targets, chemical series, and screening settings remains to be established.

## 4 Conclusions

We present the first public OpenBind release, an experimental structure–affinity dataset for enteroviral 2A protease generated through crystallographic fragment screening and follow-on optimisation. The release contains 925 crystallographic binding events from 699 compounds and affinity measurements for 601 compounds, with 490 compounds retained for the final quantitative affinity-prediction benchmark. After curation of the structural data, 881 complexes were retained, including 802 follow-on complexes used for the primary docking and cofolding benchmarks. We also constructed a virtual-screening benchmark of 2,084 follow-on compounds, comprising 566 binders and 1,518 suspected non-binders. Together, these resources support evaluation of pose prediction, receptor-state sensitivity, affinity prediction, and compound prioritisation within a coherent experimental campaign.

Our benchmark results highlight both the promise and limitations of current structure-based modelling approaches. Docking performed well when the cognate ligand-bound receptor conformation was available, but deteriorated substantially under cross-docking into less compatible receptor states. Cofolding partially mitigated this receptor-state mismatch by allowing the binding pocket to adapt during prediction, although accurate complex generation and ranking remained challenging. Fine-tuning OpenFold3-p2 on fragment-bound structures improved prediction on follow-on compounds, indicating that fragment-screening data can provide useful target-specific information. Affinity prediction remained challenging, with simple physicochemical baselines competitive with several structure-based models.

These tasks test different capabilities and should not be reduced to a single ranking of model quality. The results are better viewed as reference baselines that expose limitations in receptor-state modelling, ranking of generated complexes, and affinity prediction beyond simple physicochemical trends. The dataset should likewise be interpreted as experimental campaign data rather than as a perfectly balanced benchmark. Its partially incomplete measurements, locally dense chemical series, receptor variability, and fragment-to-follow-on progression reflect realistic discovery practice and help reveal how and why methods fail. Because these results derive from a single protein target and one target-focused discovery campaign, the relative performance of individual methods may depend on the specific receptor dynamics and chemical series represented here. Evaluation across additional targets will therefore be required to establish how broadly the conclusions generalise.

More broadly, this release demonstrates the OpenBind data-generation model: producing open, target-focused datasets that connect crystallographic structures, affinity measurements, and medicinal chemistry progression. Future releases across additional targets will extend this framework for evaluating model generalisation, target-specific adaptation, and structure-based AI methods aligned with practical molecular discovery.

## Conflicts of interest

EJG is a Director of MedChemica Ltd, MedChemica Consultancy Ltd, and MedChemica Holdings Ltd, and is a shareholder in MedChemica Holdings Ltd and AstraZeneca plc. JDC is a current member of the Scientific Advisory Board of OpenEye Scientific Software and has equity in and serves as the Chief Executive Officer of Achira, Inc., which is engaged in the creation of open foundation simulation models for drug discovery. CMD is a member of the Scientific Advisory Board of Fusion Antibodies plc and AI Proteins, and is a founder of DaltonTx. MA is a member of the scientific advisory boards of Cyrus Biotechnology, Deep Forest Sciences, Nabla Bio, Oracle Therapeutics, and Achira. The remaining authors declare no competing interests.

## Data and Code Availability

**Experimental data:** Experimental structures and compound data are available through Fragalysis at https://fragalysis.diamond.ac.uk/viewer/react/preview/target/A71EV2A/tas/lb42888-1 and are archived on Zenodo at https://zenodo.org/records/20026661.

**Prepared and benchmark data:** The prepared dataset, docking outputs, and cofolding predictions used in this study are archived on Zenodo at https://zenodo.org/records/20798528.

**GitHub:** Benchmarking and plotting code is available at https://github.com/OpenBind-Consortium/EV-A71_2A_benchmark.

## Acknowledgements

We thank everyone involved in generating and processing data, and preparing this release, as well as the funders supporting OpenBind.

OpenBind received funding from the UK Department of Science, Innovation and Technology under grant number G2-SCH-2025-06-16537. Research reported here was supported in part by NIAID of the National Institutes of Health under award number U19AI171399. The authors acknowledge the use of resources provided by the Isambard-AI National AI Research Resource (AIRR). Isambard-AI is operated by the University of Bristol and is funded by the UK Government’s Department for Science, Innovation and Technology (DSIT) via UK Research and Innovation; and the Science and Technology Facilities Council [ST/AIRR/I-A-I/1023]. The authors also acknowledge Diamond Light Source for beamtime on beamlines I03 and I04-1 under proposal LB32627. JDC acknowledges support from NIH grant P30 CA008748 and the Sloan Kettering Institute.

Disclaimer: the content is solely the responsibility of the authors and does not necessarily represent the official views of the National Institutes of Health.

## Notes

https://zenodo.org/records/20026661

https://zenodo.org/records/20798528

https://github.com/OpenBind-Consortium/EV-A71_2A_benchmark

