## Supplementary Information for "The first OpenBind release: An open experimental structure–affinity dataset and benchmark for structure-based AI"

[1]Department of Statistics, University of Oxford, Oxford OX1 3LB, United Kingdom
[2]Department of Systems Biology, Columbia University, New York, NY, United States
[3]Diamond Light Source, Harwell Science and Innovation Campus, Didcot OX11 0DE, United Kingdom
[4]Research Complex at Harwell, Harwell Science and Innovation Campus, OX11 0FA, Didcot, United Kingdom
[5]MedChemica Consultancy Ltd, Macclesfield, Cheshire, SK11 6DU, United Kingdom
[6]Computational and Systems Biology Program, Sloan Kettering Institute, Memorial Sloan Kettering Cancer Center, New York, NY, United States
[7]Centre for Medicines Discovery, University of Oxford, Old Road Campus, Roosevelt Drive, OX3 7DQ, Headington, United Kingdom
[8]Department of Biochemistry, University of Johannesburg, Auckland Park 2006, South Africa
[9]OpenBind, https://openbind.uk/

### S1 Supplementary methods

#### S1.1 Experimental data collection details

##### Protein production and purification

All crystallographic and biophysical experiments used CVA16 2A protease strain G10 as a close experimental surrogate for EV-A71 2A protease. These two proteins differ in only five positions in the amino acid sequence, all outside the catalytic site, and these substitutions are not expected to substantially alter protease activity [1]. For crystallographic experiments, recombinant CVA16 2A protease was expressed in *Escherichia coli* BL21(DE3)-RR cells as an N-terminal $His_6$-SUMO fusion protein and purified using affinity and size-exclusion chromatography according to a previously described protocol [2].

For affinity measurements, a separate biotinylated CVA16 2A protease construct was produced to enable streptavidin-mediated immobilisation on Creoptix sensor chips. Expression, purification, and biotinylation procedures followed the corresponding large-scale production protocol [3].

##### Crystallisation and structure determination

Crystallisation experiments were performed using CVA16 2A protease following previously described procedures developed for high-throughput fragment screening and ligand-soaking campaigns [4]. In the subsequent XChem crystallographic fragment screen [5], fragments from the DSi-Poised [6], SpotXplorer [7], FragLites [8], PepLites [9], York3D [10], MiniFrags [11], and Covalent MiniFrags libraries [12] were soaked into pre-formed crystals at final concentrations of approximately 50–100 mM. Together, these libraries were selected to sample a diverse range of fragment chemotypes and interaction modalities, spanning very small fragments, three-dimensional aliphatic ring systems, pharmacophore-focused fragments, and covalent fragments. Follow-on compounds generated during subsequent optimisation campaigns were soaked at nominal concentrations of 2–10 mM [1].

Crystals were harvested, cryo-cooled, and measured at the macromolecular crystallography beamlines I03 and I04-1 at Diamond Light Source. Diffraction datasets were processed using automated data-processing pipelines at Diamond and analysed through the XChemExplorer framework [13]. Initial map generation was performed using DIMPLE [14], and ligand restraints were generated using GRADE or GRADE2 [15]. Ligand-binding events were identified using PanDDA2 [16] where statistical background-map analysis was required. For later follow-on datasets in which ligands were clearly visible in conventional $2F_o - F_c$ electron-density maps, ligands were identified and modelled directly without PanDDA2. Iterative model building and refinement were performed using Coot [17] together with REFMAC [18] or BUSTER [19] through the XChemExplorer platform.

##### Affinity measurements

Binding affinities were measured using grating-coupled interferometry (GCI) on a Creoptix WAVEsystem employing waveRAPID kinetics [20]. To establish robust assay conditions, a multi-stage design-of-experiments workflow was performed to evaluate protein stability, immobilisation strategies, buffer composition, pH, detergent conditions, and long-term assay stability. Detailed experimental procedures are provided in the associated protocols.io workflow [21].

A biotinylated CVA16 2A protease construct was immobilised through streptavidin–biotin capture chemistry. Production measurements were performed using a running buffer consisting of 10 mM HEPES pH 7.0, 100 mM NaCl, 0.005% (v/v) Tween-20, 0.5 mM TCEP, and 2% (v/v) DMSO, which supported stable performance during extended assay validation experiments.

Sensorgrams were analysed using the standard Creoptix software workflow and manually reviewed by trained experimentalists. Association rates ($k_a$), dissociation rates ($k_d$), equilibrium dissociation constants ($K_D$), and associated fit-quality metrics were extracted using a 1:1 binding model. Measurements failing predefined quality-control criteria were excluded from downstream analyses.

#### S1.2 Dataset curation and benchmark construction

##### S1.2.1 Affinity benchmark construction

###### Affinity data filtering

Affinity measurements were filtered at the individual measurement level before construction of the compound-level benchmark. Only sample measurements were retained, while control measurements were excluded. Measurements were removed when the fitted association rate constant, $k_a$, reached its fitting boundary of 100, the dissociation rate constant was equal to $10^{-7}$, or $R_{\max}$ was equal to 0.001, as these values indicate that the corresponding parameter had reached a fitting boundary.

Measurements were additionally excluded when the reported relative error in either $k_a$ or $k_d$ exceeded 200%, when $\sqrt{\chi^2}$ exceeded 2, or when $R_{\max}$ was below 0.5. Compounds for which only a single qualifying measurement remained after filtering were retained. Application of these criteria reduced the affinity dataset from 601 measured compounds to 494 compounds with qualifying affinity measurements.

###### Compound-level aggregation

Benchmark evaluation was performed at the compound level rather than the crystallographic binding-event level. Where multiple crystallographic structures or corresponding model predictions were available for the same compound, predicted affinities were averaged. This yielded one predicted affinity and one experimental affinity for each of the 494 compounds retained after affinity filtering.

Binding events were excluded from affinity-prediction analyses if they were considered suspected artefacts or failed PoseBusters validity checks. After applying these exclusions at the binding-event level and consolidating predictions at the compound level, four compounds were removed from the benchmark, yielding a final affinity-prediction set of 490 compounds.

##### S1.2.2 Virtual screening benchmark construction and characterisation

###### Benchmark construction

We constructed a virtual screening benchmark by combining curated crystallographic binders with suspected non-binders from the soaking campaign. Binders were defined as compounds for which an experimental ligand-bound crystal structure was obtained. Protein–ligand complexes that were not PoseBusters-valid or were flagged for potential crystal-packing effects were excluded from the benchmark binder set, consistent with the structural benchmark eligibility criteria.

Suspected non-binders were selected from compounds for which no bound crystal structure was observed. To reduce ambiguity in this negative set, we excluded compounds with any reported LigandCC value and retained only experiments with sufficient crystallographic data quality to support suspected non-binder assignment. These experiments were required to have mounted and diffracted crystals, the expected C121 or C2 space group, resolution $\leq 2.75$ Å, successful DIMPLE processing, and $R_{\text{free}} < 0.4$.

Before benchmark assembly, compound structures were canonicalised and deduplicated at the compound level. Duplicate records corresponding to the same canonical isomeric SMILES were collapsed to a single benchmark entry. Where the same compound appeared in both binding and non-binding records, binder status was retained if the compound had been observed to bind in any qualifying experiment. Records with ambiguous compound identifiers or structures for which a three-dimensional conformer could not be generated with RDKit were excluded.

Binders retained their experimental fragment-screen annotation. Suspected non-binders originating from designated fragment libraries were classified as fragments. Compounds from the designed compound collections were classified as fragment-like when their molecular weight was $< 250$ Da or their heavy atom count was $\leq 18$; only compounds exceeding both thresholds were kept as follow-on compounds.

The benchmark was restricted to follow-on compounds. Curated binders originating from the fragment screen were excluded. Suspected non-binders originating from designated fragment libraries were also excluded. In addition, compounds from the designed compound collections were excluded when their molecular weight was below 250 Da or their heavy-atom count was at most 18, in order to remove smaller, fragment-like compounds. The final virtual screening benchmark contained $n = 2{,}084$ compounds, comprising 566 binders and 1,518 suspected non-binders.

**Benchmark characterisation**

The chemical composition of the virtual screening benchmark was characterised using ECFP4 fingerprints and physicochemical descriptors. Compounds were represented using 2,048-bit ECFP4 fingerprints with chirality enabled. Pairwise Tanimoto similarities were calculated to identify the nearest binder and nearest suspected non-binder for each compound, and the fingerprint matrix was projected into two dimensions using UMAP with the Jaccard distance metric, 30 nearest neighbours, a minimum distance of 0.1, and a random seed of 1. Physicochemical descriptors calculated using RDKit included molecular weight, heavy-atom count, cLogP, topological polar surface area, numbers of hydrogen-bond donors and acceptors, rotatable-bond count, aromatic-ring count, and fraction of $sp^3$-hybridised carbons.

#### S1.3 Computational benchmarking and analysis

##### S1.3.1 Molecular docking calculations

**Ligand and receptor preparation for docking**

Ligand and receptor preparation was standardised across all docking settings. Ligand preparation started from the crystallographic ligand structures provided in SDF format. Ligands were protonated at pH 7.4 using OpenBabel 3.1.0 [22] and subsequently processed with RDKit 2025.03.6 [23]. After sanitisation, five starting conformers were generated for each compound using the ETKDGv3 algorithm [24] with a fixed random seed. The resulting conformers were minimised using the UFF force field [25] and written as SDF files for docking.

Receptor preparation started from ligand-free crystallographic receptor structures in which bound ligands, solvent molecules, and cofactors had been removed. Receptors were prepared using PDBFixer v1.12 [26], which was used to resolve alternate conformations and add hydrogens at pH 7.4. Because enteroviral 2A protease is a monomeric enzyme, just a single protein chain was retained. For structures containing multiple protein chains, only the chain with the largest number of residues within 6 Å of the crystallographic ligand was retained.

The experimentally determined apo structure (PDB ID: 8POA) and AlphaFold2-predicted apo model were processed using the same PDBFixer-based preparation protocol as the crystallographic receptors. Apo and fragment-informed receptor structures were aligned to the coordinate frame of each target crystallographic receptor using ProDy v2.6.1 [27], enabling a common binding-site definition to be used across receptor choices.

For fragment-informed cross-docking, ECFP4 fingerprints [28] were generated in RDKit as Morgan fingerprints with radius 2 and 2048 bits. Each follow-on compound was paired with the fragment-screen compound with the highest Tanimoto similarity, and the corresponding fragment-bound receptor structure was used for docking.

**Docking parameters**

Docking was performed using GNINA v1.3.2 [29], smina [30] with the Vina v1.1.2 scoring function [31], and DiffDock-L v1.1.3 [32]. GNINA and smina used the prepared receptor structures in PDB format and the five prepared ligand conformers in SDF format. Each conformer was docked independently using a cubic search box with side length 25 Å, centred on the centroid of the crystallographic ligand.

GNINA was run with the default ensemble CNN model, exhaustiveness 8, and up to eight output poses per conformer. smina was run with the Vina scoring function, exhaustiveness 8, and up to eight output poses per conformer. Consistent random seeds were used for both methods. This yielded up to 40 poses per complex and receptor scenario for each method. Poses generated from the five starting conformers were pooled, reranked using the corresponding method-specific score, and the 25 highest-ranked poses were retained for evaluation.

DiffDock was run using the same prepared receptor and ligand inputs. For each complex and receptor scenario, five independent inference runs were performed using the default settings. Although seed labels were used to organise the output directories, DiffDock did not allow the random seed to be set explicitly. Each run generated 40 poses with associated confidence scores. The five highest-confidence poses from each run were pooled, reranked by DiffDock confidence, and the top 25 poses were retained for evaluation.

##### S1.3.2 Virtual-screening benchmark calculations

###### Docking-based virtual screening

Docking poses were generated using the fragment-informed GNINA and smina protocols described above. For each compound, the receptor structure was selected from the curated fragment-bound complexes using the highest ECFP4 Tanimoto similarity. The docking box was centred on the crystallographic ligand of the selected fragment-bound complex. For each compound, the highest-ranked pose was retained according to the corresponding docking scoring function.

###### Cofolding-based virtual screening

For the cofolding benchmark, the predicted complex with the highest pairwise interface predicted template modelling score (pair ipTM) was retained for each compound. Compounds were then ranked using the pair ipTM value of the selected complex. Pair ipTM reflects model confidence in the predicted protein–ligand interface and was used here directly as the cofolding-based virtual-screening score.

###### Baselines and performance assessment

Molecular weight and cLogP were calculated for all compounds using RDKit and included as simple physicochemical baselines. Scores were oriented such that larger values corresponded to higher predicted binding likelihood. For all scoring approaches and baselines, compounds were ranked according to the corresponding score and evaluated on the common set of compounds available across all methods. Virtual-screening performance was quantified using ROC-AUC, average precision, and enrichment factors at 1%, 2%, and 5%. ROC-AUC and average precision were calculated using scikit-learn, while enrichment factors were calculated using NumPy by comparing the binder fraction within the top-ranked 1%, 2%, and 5% of compounds with the overall binder prevalence in the evaluation set.

#### S1.4 OpenFold3 fine-tuning

OpenFold3-p2 was fine-tuned on the 79 curated fragment-bound structures from the OpenBind enteroviral 2A protease dataset and evaluated on the follow-on compound benchmark. For structures containing two protein copies in the asymmetric unit, only the chain associated with the modelled ligand-binding event was retained. The resulting single-chain protein–ligand complexes were processed using the OpenFold data-processing pipeline, and MSAs were generated with the ColabFold MSA server [33].

We conducted a hyperparameter sweep over batch sizes $\{8, 16, 32\}$ and learning rates $\{10^{-2}, 10^{-3}, 10^{-5}, 10^{-6}\}$. The learning rate was linearly warmed up for 100 steps. Model performance showed little sensitivity to batch size, while higher learning rates introduced training instabilities. Based on these results, we selected a batch size of 8 to minimise computational cost, paired with a learning rate of $10^{-3}$, which offered the best trade-off between training stability and convergence speed. We trained for 1,000 steps ( 8 hours wall-clock time), by which point the protein–ligand LDDT metrics had converged. Each fine-tuning experiment was performed using 8 NVIDIA GH200 GPUs.

### S2 Supplementary analyses

#### S2.1 Similarity to public structural data

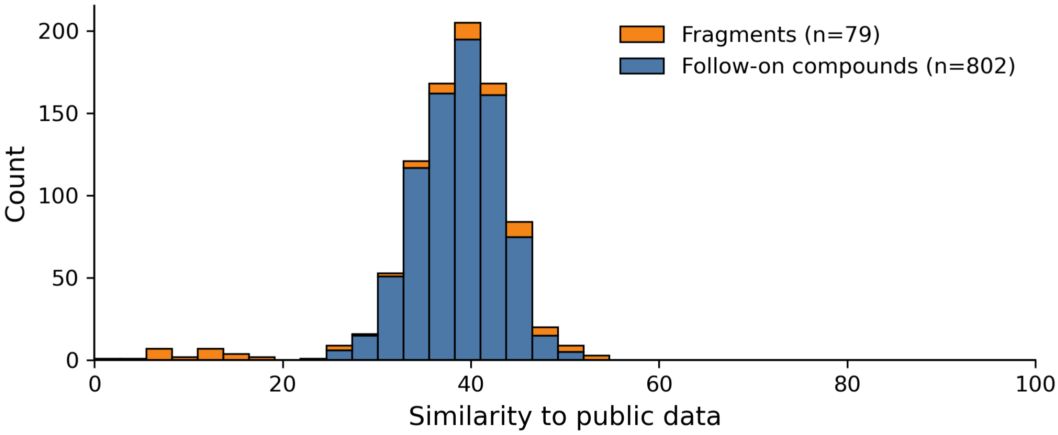

Figure S1: Distribution of ligand–pocket similarity scores between the 881 curated OpenBind enteroviral 2A protease complexes and public protein–ligand structures released on or before 1 June 2023, corresponding to the reported Boltz-2 training cutoff. Fragment-bound ($n = 79$) and follow-on complexes ($n = 802$) are shown in orange and blue, respectively. Similarity was calculated using the Runs N' Poses ligand–pocket metric.

Table S1: Summary statistics for ligand–pocket similarity scores between curated OpenBind enteroviral 2A protease complexes and public protein–ligand structures. Values outside parentheses were calculated using the AlphaFold3 reference cutoff of 30 September 2021, while values in parentheses use the Boltz-2 cutoff of 1 June 2023 where the result differed. Statistics are reported for all 881 curated complexes and separately for the 79 curated fragment-bound and 802 curated follow-on complexes. Threshold rows report the percentage of complexes with similarity scores below the indicated value.

| Metric | All curated complexes | Fragment-bound complexes | Follow-on complexes |
|---|---|---|---|
| $n$ | 881 | 79 | 802 |
| Min. | 0.00 | 0.00 | 25.43 |
| Q1 | 35.51 | 14.96 | 35.89 |
| Median | 38.85 | 37.65 | 38.94 |
| Q3 | 42.00 | 43.94 | 41.90 |
| IQR | 6.48 | 28.98 | 6.01 |
| Mean | 38.14 | 31.73 (31.76) | 38.77 |
| Max. | 54.68 | 54.68 | 51.66 |
| $< 30$ (%) | 5.68 | 36.71 | 2.62 |
| $< 40$ (%) | 59.59 | 60.76 | 59.48 |
| $< 50$ (%) | 99.32 | 96.20 | 99.63 |

#### S2.2 Relationship between fragment hits and follow-on compounds

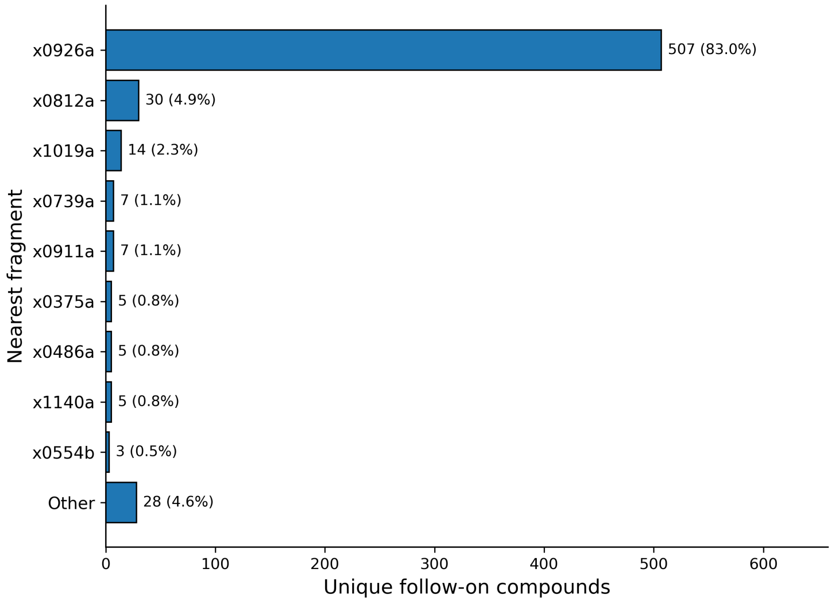

Figure S2: Nearest-fragment assignments for the 611 unique curated follow-on compounds. Each follow-on compound was assigned to the curated fragment with the highest ECFP4 Tanimoto similarity among all 66 unique curated fragment-screen compounds. The most frequently assigned fragments are shown individually, with all remaining fragments grouped as “Other”. Labels report the number and percentage of follow-on compounds assigned to each fragment.

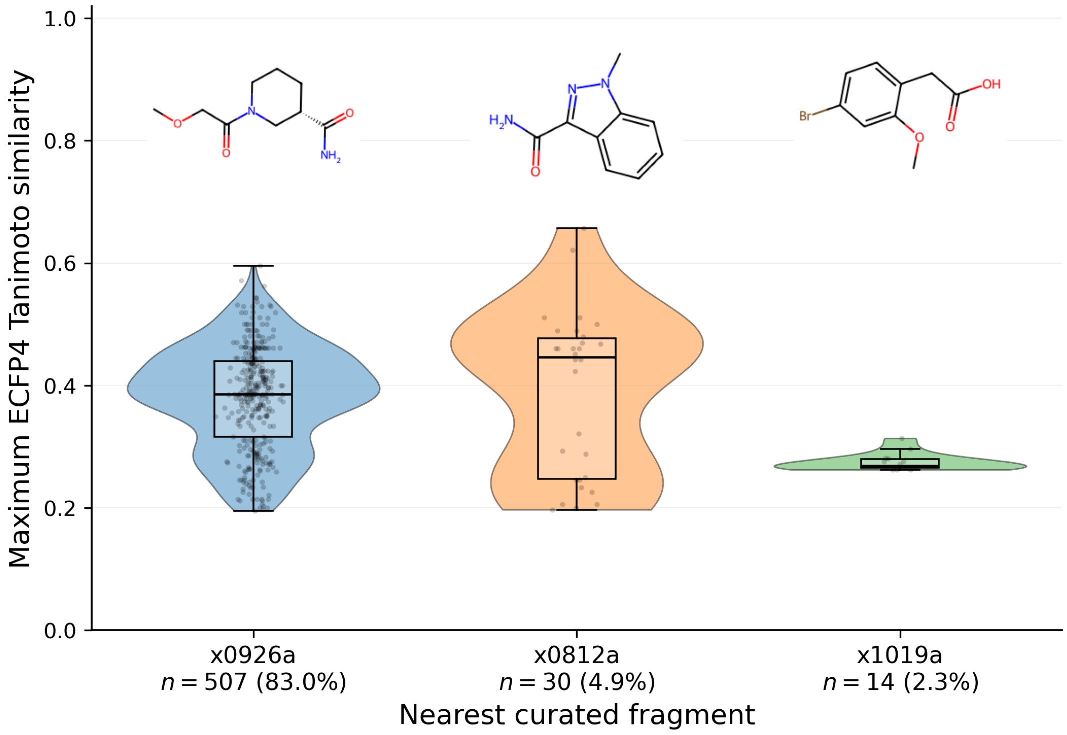

Figure S3: Distribution of maximum ECFP4 Tanimoto similarities for follow-on compounds in the three largest nearest-fragment groups. Each value is the maximum similarity of a unique curated follow-on compound to any of the 66 unique curated fragment-screen compounds. The corresponding fragment structure is shown above each violin plot. Group sizes indicate the number of follow-on compounds assigned to each nearest curated fragment.

#### S2.3 Multi-structure fragment cross-docking

To assess whether the use of multiple experimentally observed pocket conformations improves cross-docking performance, we cumulatively combined predictions generated against the five most similar fragments according to ECFP4 Tanimoto similarity.

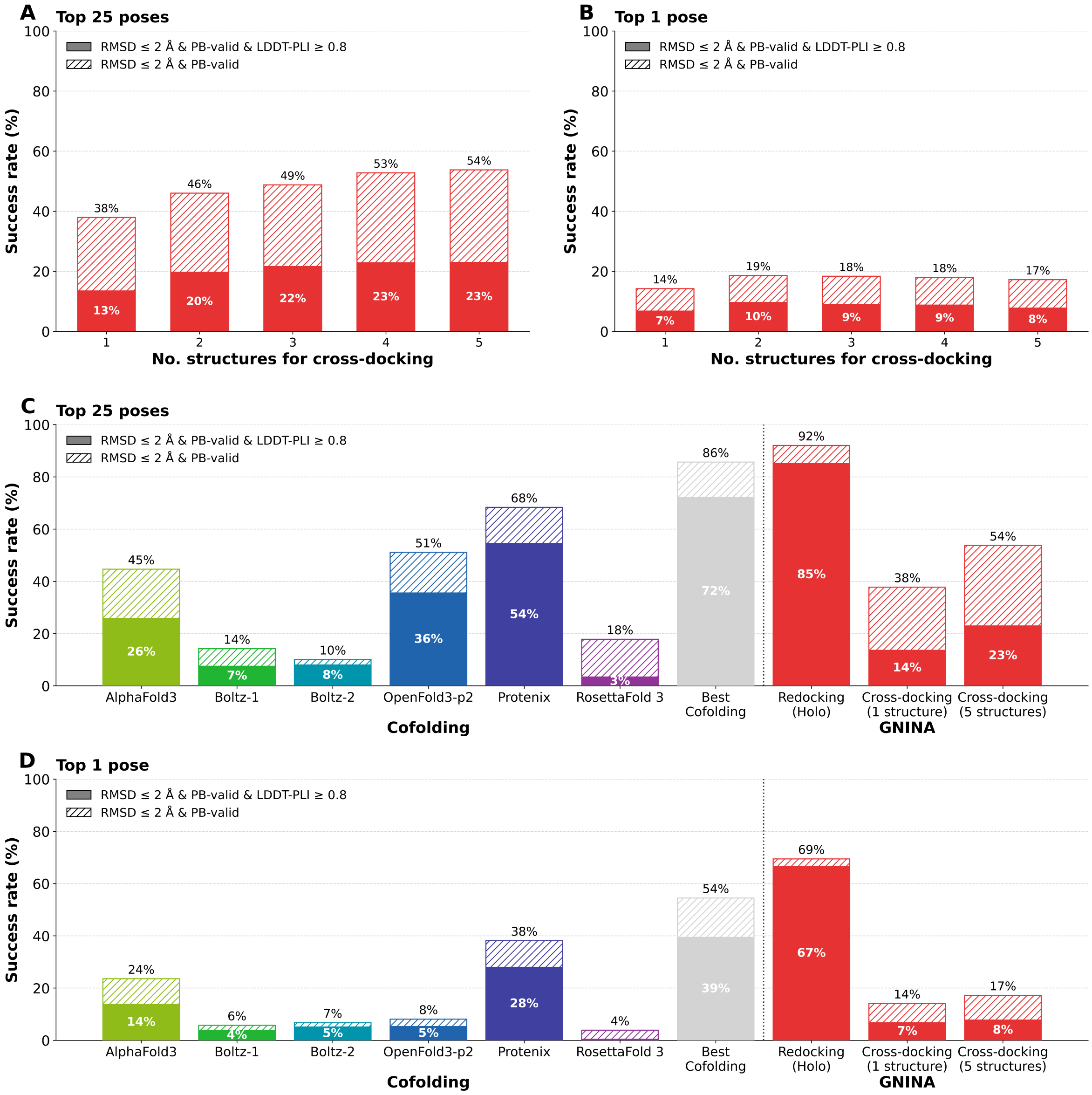

Figure S4: Multi-receptor fragment-informed cross-docking performance on the curated OpenBind enteroviral 2A protease follow-on benchmark ($n$ = 802 binding events). Panels **A** and **B** show GNINA Top-25 and Top-1 performance, respectively, after cumulatively pooling and reranking predictions generated using the one to five most chemically similar fragment-bound receptor structures. Panels **C** and **D** compare the corresponding five-receptor results with cofolding methods, single-receptor fragment-informed cross-docking, and GNINA redocking. Solid bars indicate predictions satisfying ligand heavy-atom RMSD $\leq 2$ Å, LDDT-PLI $\geq 0.8$, and PoseBusters validity; hatched extensions indicate additional PoseBusters-valid predictions satisfying the RMSD criterion only.

#### S2.4 Failure mode analysis for cofolding methods

Traditional docking approaches typically place ligands on the surface of a rigid receptor structure, typically within a predefined volume centred on a binding site of interest. In contrast, cofolding methods must model the receptor conformation and ligand binding mode simultaneously. As a result, cofolding predictions can be inaccurate not only due to incorrect ligand placement, but also because of incorrect receptor conformations.

To assess the underlying causes of inaccurate cofolding predictions, a failure mode analysis was conducted to compare sources of failed predictions across all benchmarked cofolding models.

**Failure mode definitions**

Although other failure modes are likely to exist, this analysis focuses on pocket identification and receptor-pocket modelling as two major contributors to cofolding failures.

To determine whether the ligand is placed within the correct binding site in a given cofolding model, the pocket recall metric is calculated. In this work, pocket recall is defined as the fraction of ground-truth receptor residues within 6.0 Å of the ligand that are also present within 6.0 Å of the ligand in the cofolded model. For a given cofolded structure, if the pocket recall $< 0.35$, the ligand is considered to be placed outside of the binding pocket, and the structure is marked as a pocket finding failure.

To assess whether the conformation of the binding pocket was correctly modelled, ligand pocket LDDT (LDDT-LP) is calculated for each cofolded structure. If the LDDT value is $< 0.8$ the pocket conformation is considered incorrect, and the structure is marked as a pocket conformation failure.

**Failure mode results**

For each cofolding method, we label the failure modes for the "best of" model for the subset of binding events in the structural benchmark set that included follow-on compounds. Next, we consider four failure modes that denote if the ligand was placed in the binding pocket, with or without the correct pocket conformation (Figure S5).

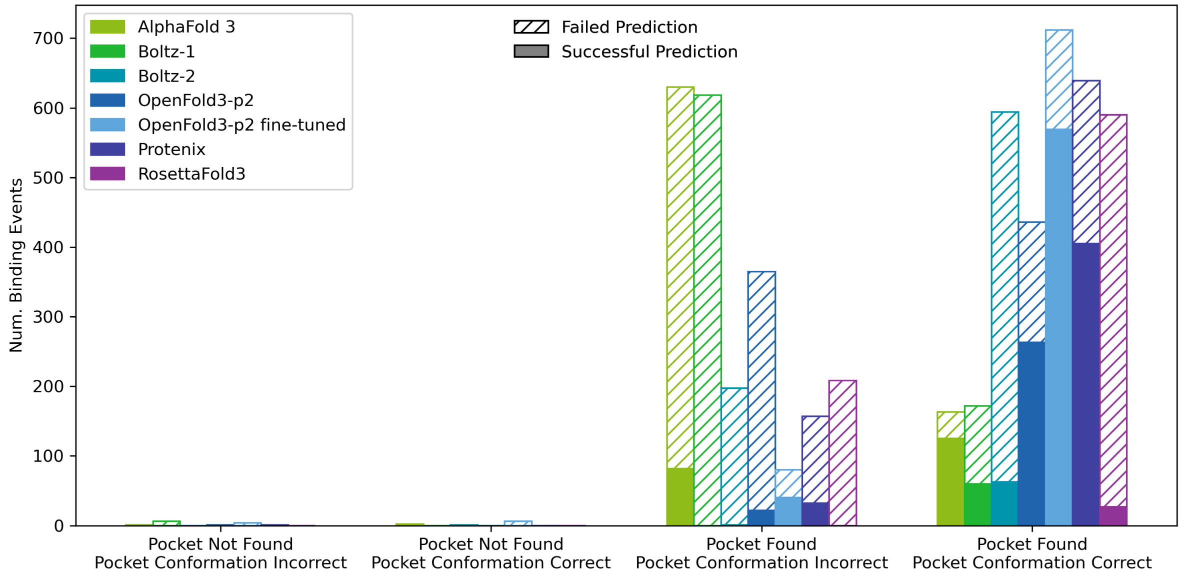

Figure S5: Number of successful and failed structural predictions for follow-on compounds in the structural benchmark set for various failure modes. For each cofolding method, the "best of" model was selected for each binding event. A model is considered to be a successful prediction if ligand RMSD $\leq 2$ Å, LDDT-PLI $\geq 0.8$, and is PoseBusters valid.

Based on this classification scheme, we observe that different cofolding methods are impacted by each failure mode with differing levels of severity. In general, ligands in the structural benchmark set are rarely placed outside of the correct binding pocket. For AlphaFold3 and Boltz-1, the primary failure mode appears to be generating incorrect conformations of the binding pocket (Figure S5).

In contrast, pocket conformational modelling is less of an issue for Boltz-2, OpenFold3-p2, Protenix, and RosettaFold3. However, when the binding pocket is modelled accurately, a large proportion of Boltz-2 and RosettaFold3 predictions fail to produce an accurate structure. This is indicative of the fact that ligand placement can remain a challenge for some cofolding models even if an accurate pocket conformation is generated.

Notably, fine-tuning OpenFold3-p2 on fragment structures reduces the proportion of predictions for which pocket-conformation failures are observed. Furthermore, the success rate improves when the ligand is placed within the correct pocket with an accurate conformation, indicating that training on fragment-screening data can alleviate issues with generating accurate ligand poses when the correct receptor structure is obtained. This result is promising, as it suggests that fine-tuning the model on an ensemble of fragment-bound structures can steer OpenFold3-p2 towards learning more accurate receptor conformations.

#### S2.5 Affinity benchmark

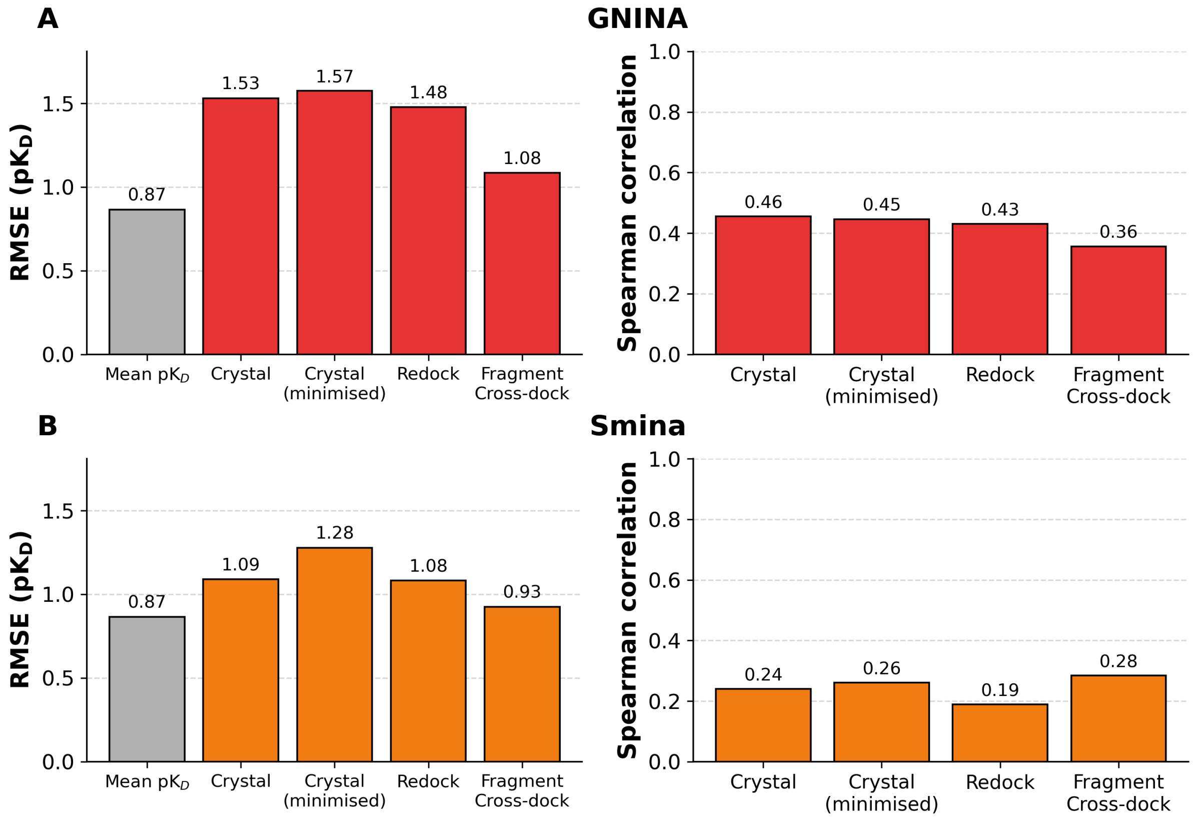

Figure S6: Affinity-prediction performance of GNINA and smina across different structural input scenarios on the final compound-level benchmark ($n = 490$). Root-mean-square error (RMSE; lower is better) and Spearman rank correlation (higher is better) are shown for GNINA (**A**) and smina (**B**). Scores were evaluated using the experimental crystal structures, the same crystal structures after local minimisation, redocked poses, and fragment-informed cross-docked poses. The RMSE obtained by predicting the mean experimental p$K_D$ for every compound is included as a reference baseline. Across all structural scenarios, GNINA showed consistently stronger rank correlation than smina, whereas smina yielded lower RMSE values.

#### S2.6 Virtual screening benchmark

##### S2.6.1 Benchmark characteristics

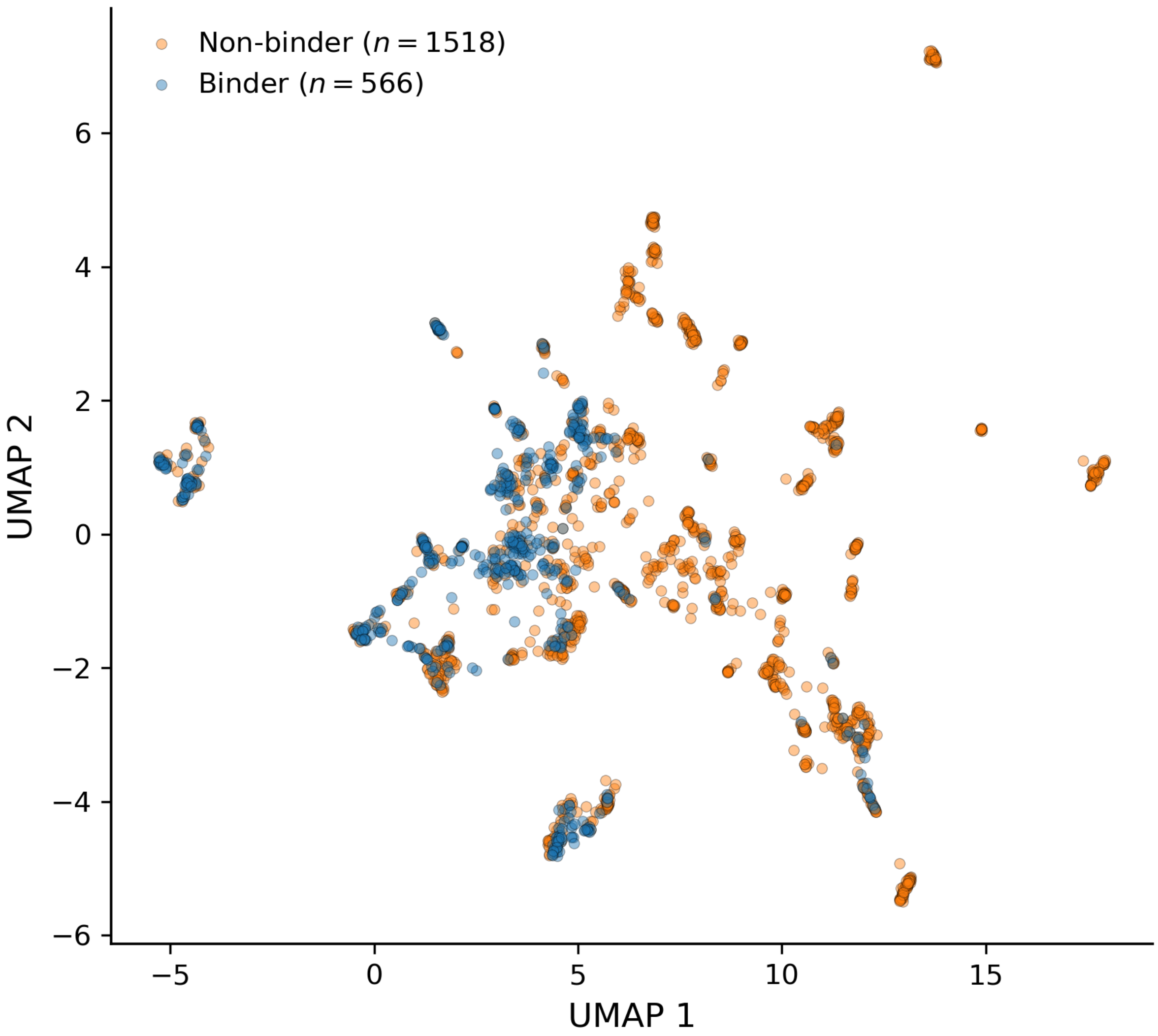

Figure S7: Chemical-space distribution of the virtual-screening benchmark ($n$ = 2,084 compounds). ECFP4 fingerprints were embedded using UMAP and coloured by crystallographic classification, with binders ($n$ = 566) shown in blue and suspected non-binders ($n$ = 1,518) in orange. Binders largely occupy regions that also contain suspected non-binders, while the broader suspected non-binder set extends into additional regions with comparatively few confirmed binders.

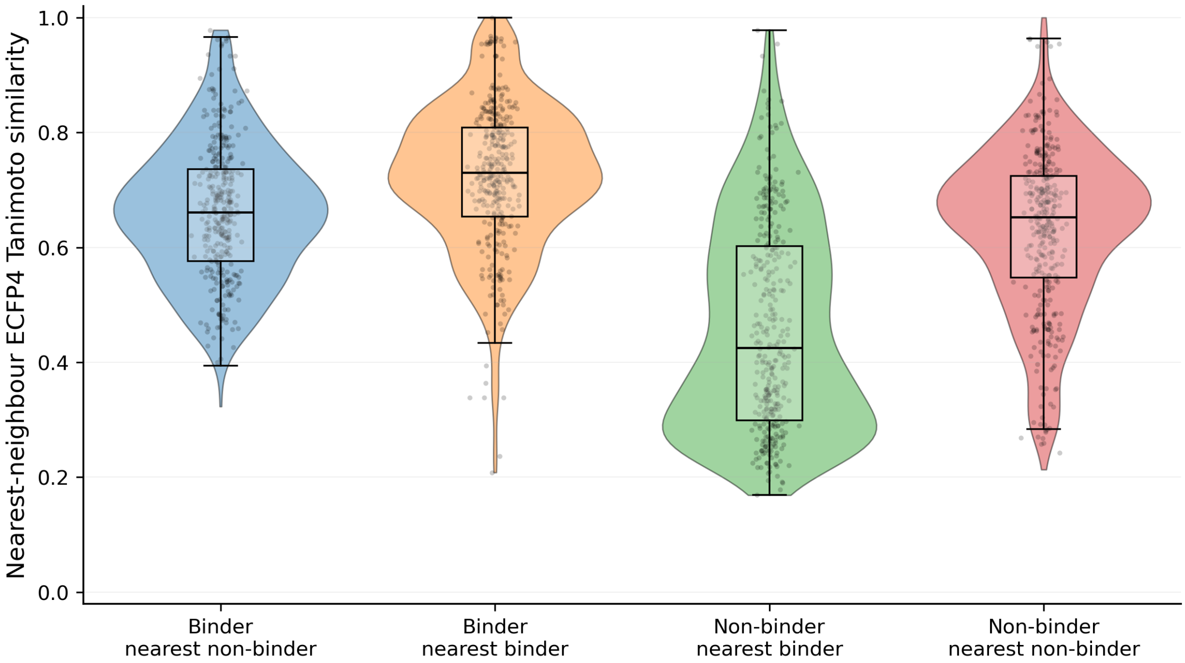

Figure S8: Nearest-neighbour chemical similarity distributions for the virtual screening benchmark. ECFP4 Tanimoto similarities were calculated from each curated binder ($n = 566$) and suspected non-binder ($n = 1{,}518$) to its nearest compound in the same and opposite class. The median similarities were 0.661 for binders to their nearest suspected non-binder, 0.730 for binders to their nearest binder, 0.425 for suspected non-binders to their nearest binder, and 0.653 for suspected non-binders to their nearest suspected non-binder.

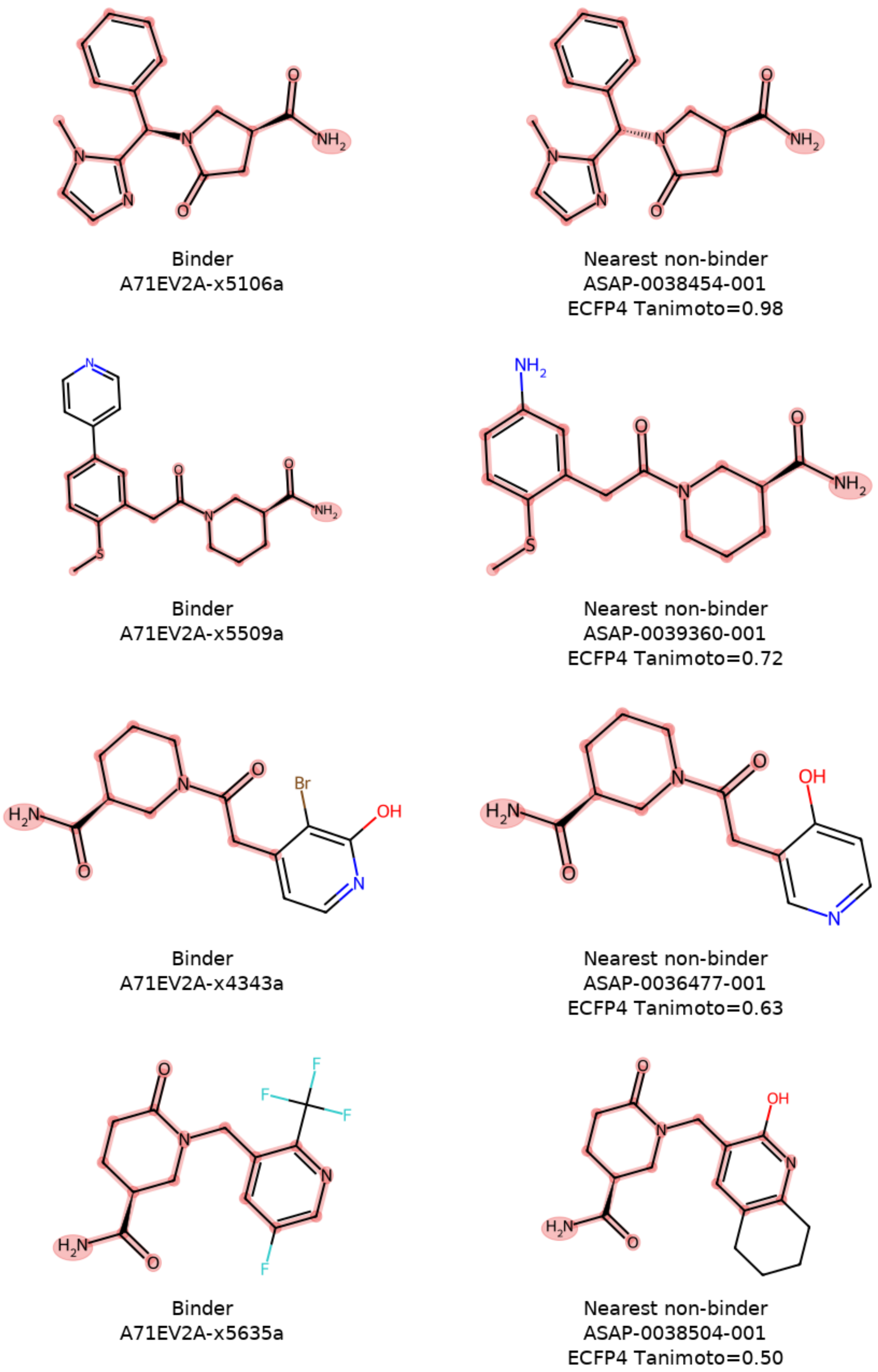

Figure S9: Examples from the virtual screening benchmark showing curated binders and their nearest suspected non-binders according to ECFP4 Tanimoto similarity. Pairs were selected at evenly spaced ranks across the similarity range of 0.5–1.0, which contains 91.0% of binder–nearest-non-binder relationships in the follow-on benchmark. Shared molecular features are highlighted in red, and the corresponding nearest-neighbour similarity is reported below each suspected non-binder. The examples illustrate that compounds with different experimental classifications can retain substantial structural similarity and, in some cases, differ only through relatively local chemical modifications.

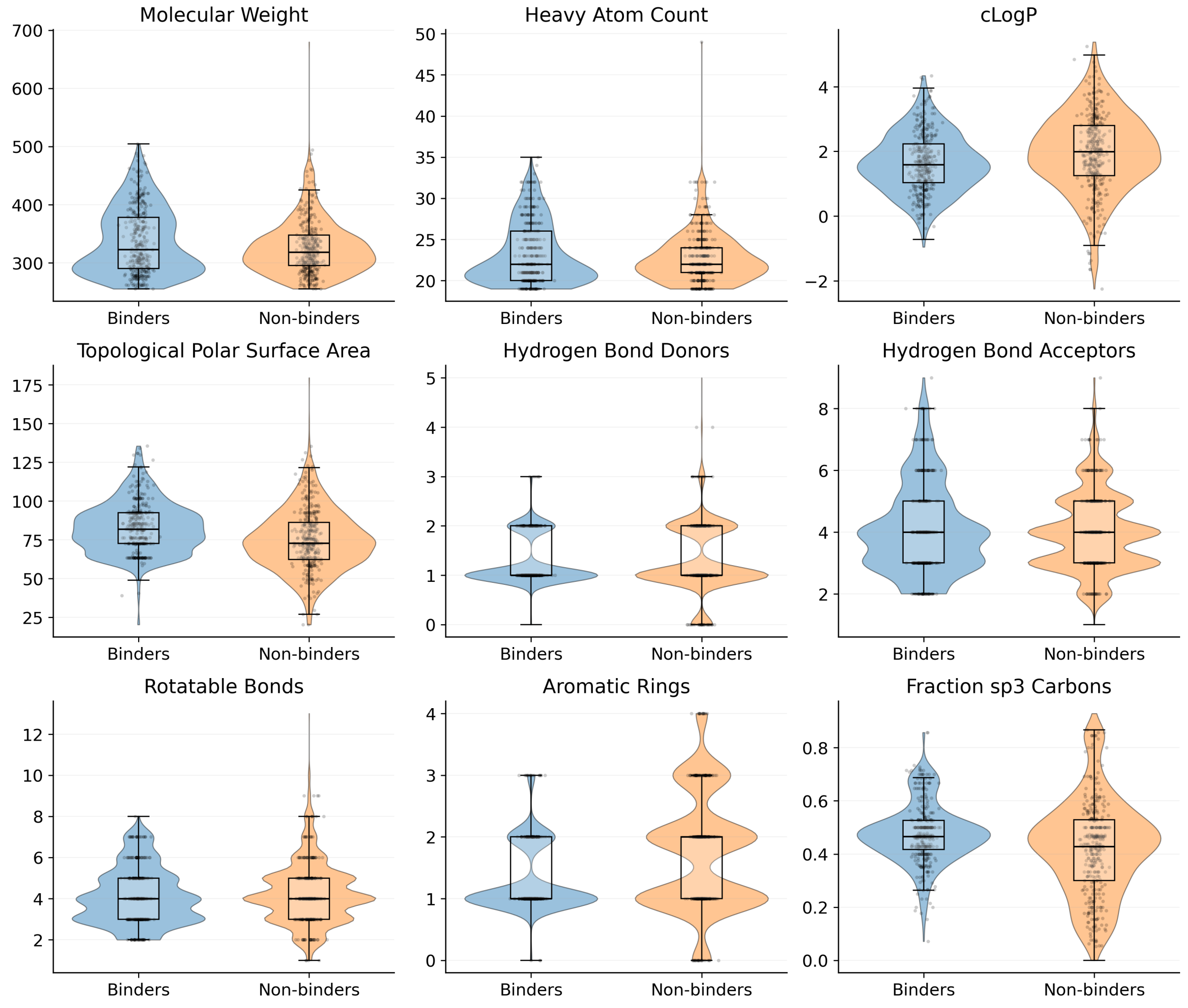

Figure S10: Physicochemical descriptor distributions for the virtual screening benchmark. RDKit descriptors were calculated for curated binders ($n$ = 566) and suspected non-binders ($n$ = 1,518). Violin plots with overlaid box plots and individual compound values show molecular weight, heavy-atom count, calculated logP (cLogP), topological polar surface area (TPSA), hydrogen-bond donor count, hydrogen-bond acceptor count, rotatable bond count, aromatic ring count, and fraction of sp$^3$-hybridised carbons (Fsp$^3$). The broadly overlapping distributions indicate that the two classes are not trivially separated by individual physicochemical descriptors.

##### S2.6.2 Virtual screening performance

Virtual screening performance was evaluated on the common benchmark set of 2,084 compounds, with 566 experimentally confirmed binders and 1,518 suspected non-binders. Performance was quantified using the area under the receiver operating characteristic curve (ROC-AUC), average precision, and enrichment factors (EFs) calculated for the top 1%, 2%, and 5% of each ranked compound list. ROC-AUC measures discrimination across the complete ranking, whereas average precision summarises binder retrieval across all ranking thresholds. Enrichment factors measure the concentration of binders among the highest-ranked compounds relative to random selection. Scores were oriented so that better predictions ranked first: native smina minimized-affinity and cLogP values were sign-inverted before evaluation so that larger final scores corresponded to higher ranking priority for all methods and baselines. For GNINA, CNN_VS was used for the primary comparison in the main text because it is specifically intended for virtual-screening ranking, while results for CNNaffinity and CNNscore are also included here for completeness.

The fine-tuned OpenFold3-p2 model showed the strongest overall performance, with the highest ROC-AUC (0.757), average precision (0.525), and EF at 5% (2.174; Table S2). Among the docking-based approaches, GNINA CNNaffinity gave the highest ROC-AUC (0.698) and average precision (0.424). GNINA CNNscore showed weaker global discrimination but stronger early enrichment, reaching an EF of 2.455 at 1% and 2.367 at 2%. The same EF at 1% was obtained by the fine-tuned OpenFold3-p2 model. GNINA CNN_VS also outperformed smina according to ROC-AUC and average precision.

The non-fine-tuned OpenFold3-p2 model performed only slightly above random according to ROC-AUC and showed limited early enrichment. The molecular-weight baseline performed only slightly better than random according to ROC-AUC and showed modest early enrichment. cLogP provided somewhat better overall discrimination than molecular weight, but its average precision remained low and its enrichment factors indicated poor retrieval among the highest-ranked compounds.

Table S2: Virtual screening performance on the common evaluation set of 2,084 compounds, comprising 566 binders and 1,518 suspected non-binders. Enrichment factors indicate binder enrichment relative to random selection within the top 1%, 2%, and 5% of the ranked compound list. For the cLogP baseline, lower cLogP values were ranked first. Higher values indicate better performance for all reported metrics.

| Method | ROC-AUC | Average precision | EF 1% | EF 2% | EF 5% |
|---|---|---|---|---|---|
| Smina | 0.624 | 0.326 | 0.526 | 0.877 | 0.912 |
| GNINA (CNN_VS) | 0.665 | 0.404 | 2.279 | 1.841 | 1.613 |
| GNINA (CNNscore) | 0.586 | 0.371 | **2.455** | **2.367** | 1.999 |
| GNINA (CNNaffinity) | 0.698 | 0.424 | 1.753 | 1.578 | 1.929 |
| OpenFold3-p2 | 0.533 | 0.279 | 1.052 | 0.877 | 0.771 |
| OpenFold3-p2-FT | **0.757** | **0.525** | **2.455** | 2.192 | **2.174** |
| Molecular weight | 0.539 | 0.345 | 1.929 | 1.666 | 1.613 |
| cLogP | 0.600 | 0.310 | 0.000 | 0.175 | 0.631 |
